# Pancreatic cancer cells breach endothelial barriers through protrusion-driven invasion or endothelial retraction

**DOI:** 10.64898/2026.09.02.749041

**Authors:** Gautier Follain, Sujan Ghimire, Monika Vaitkevičiūtė, Helene Helo, Joanna W. Pylvänäinen, Frédéric Fercoq, Aleksi Isomursu, Megan R. Chastney, Omkar Joshi, Marco De Donatis, Ermei Mäkilä, James R.W. Conway, Marko Salmi, Sara A. Wickström, Leo M. Carlin, Johanna Ivaska, Guillaume Jacquemet

## Abstract

Extravasation, the exit of circulating cancer cells from blood vessels, is a critical yet poorly understood step in metastatic dissemination. Here we show that pancreatic ductal adenocarcinoma (PDAC) cells can breach endothelial barriers through two mechanistically distinct modes of extravasation. MIA PaCa-2 cells breach endothelial junctions via filopodia-like protrusions, enabling access to and spread across the basal extracellular matrix (ECM). By contrast, AsPC-1 cells remain rounded atop the endothelium and cross the barrier by triggering rapid retraction of neighbouring endothelial cells. These distinct extravasation modes were also observed in zebrafish larvae. In the mouse lung, AsPC-1 cells arrest, survive, induce endothelial detachment from the basal lamina, and extravasate through this retraction mechanism before metastatic outgrowth. Mechanistically, AsPC-1-secreted factors are sufficient to destabilise endothelial monolayers, and AsPC-1 cells also induce endothelial apoptosis; however, blocking apoptosis does not prevent barrier disruption. By contrast, treatment with saracatinib, a Src-family kinase inhibitor, protects endothelial barriers, limits early vascular disruption in the lung, and delays metastatic outgrowth. Together, these findings reveal that PDAC cells can extravasate via mechanistically distinct routes, suggesting that effective anti-metastatic strategies may need to target multiple modes of endothelial barrier breach rather than a single pathway.

## Introduction

Metastasis accounts for most cancer-related mortality, and pancreatic ductal adenocarcinoma (PDAC) is among the deadliest solid malignancies. Despite therapeutic advances, five-year relative survival is approximately 13% overall and 3% for patients diagnosed with distant disease (1). The lungs, liver, peritoneum, and distant lymph nodes are among the most common sites of PDAC metastasis (1–3).

Metastatic dissemination is a multistep process in which cancer cells can enter the bloodstream, withstand haemodynamic and immune challenges, arrest within distant microvascular beds and cross the vascular wall. Only a small proportion of disseminated cells ultimately form metastases. There are several bottlenecks in the process, particularly post-extravasation survival and early metastatic outgrowth (4, 5). Within the microvasculature, circulating tumour cells can arrest either through occlusion or active adhesion to endothelial cells (6–12). Following arrest, dissemination typically requires transmigration across both the endothelial layer and the underlying vascular basement membrane. Once in the parenchyma, disseminated cells may die, enter dormancy or initiate metastatic outgrowth (13).

Despite its importance, the cellular mechanisms of cancer-cell extravasation remain poorly understood. Extravasation is often framed as the cancer-cell counterpart of leukocyte diapedesis, but this analogy is largely incomplete (14, 15). Leukocytes can cross the endothelium rapidly via tightly regulated paracellular or transcellular routes, thereby supporting efficient resealing of the vascular barrier. By comparison, circulating tumour cells and their nuclei are typically larger, making passage through narrow endothelial openings challenging. Cancer cells are also mechanically heterogeneous, and properties such as size, deformability and viscosity influence vascular arrest and extravasation efficiency (16).

Consistent with this diversity, several mechanisms of cancercell extravasation have been described, including protrusion-extension into the basement matrix and paracellular passage through destabilised endothelial junctions (8, 17, 18), endothelial remodelling (9, 19, 20), or induction of endothelial necroptosis (21). Together, these observations indicate that cancer-cell extravasation does not follow a single leukocyte-like programme (15). Importantly, mechanistic studies (particularly those employing microfluidic vascular models) have relied disproportionately on breast cancer cells, especially MDAMB-231 cells, leaving open the possibility that other cancer types may use other modes of extravasation (22).

Here, we combine complementary *in vitro* and *in vivo* imaging approaches to show that PDAC cells can breach endothelial barriers through at least two distinct extravasation modes. MIA PaCa-2 cells use protrusion-led extravasation, extending filopodia-like, actin-rich structures toward endothelial junctions and through junctional discontinuities before spreading on the underlying basal extracellular matrix. In contrast, AsPC-1 cells remain rounded atop endothelial cells, inducing retraction and detachment of the basal lamina and breaching the endothelial barrier. This endothelial retraction mode of extravasation involves cancer-cell-secreted factors that destabilise endothelial monolayers. Although endothelial apoptosis accompanies barrier failure, it is not required. Instead, treatment with saracatinib, an inhibitor of Src-family kinases, protects endothelial barriers, limits early vascular disruption during lung extravasation, and delays metastatic outgrowth. Together, these findings reveal mechanistic heterogeneity in PDAC extravasation and identify AsPC-1-induced endothelial destabilisation as a targetable vulnerability in metastatic colonisation.

## Results

### PDAC cells breach endothelial barriers during transmigration

We recently reported that PDAC cells can arrest on endothelial monolayers as efficiently as leukocytes under flow *in vitro* (12). However, arrest is only the first step of extravasation, and it remains unclear whether PDAC cells subsequently cross the endothelium through processes resembling leukocyte diapedesis.

To investigate how PDAC cells cross endothelial barriers, we first compared their behaviour with that of primary neutrophils in an *in vitro* perfusion assay. Under flow, primary neutrophils transmigrated within minutes and rapidly moved underneath the endothelial monolayer without detectable loss of barrier integrity (Fig. 1A), consistent with previous studies (23, 24). In contrast, under identical conditions, PDAC cells remained adherent atop the endothelium and completed transmigration in hours rather than minutes (Fig. 1A and Fig. 1B). This pronounced kinetic difference suggests that PDAC cells do not use a leukocyte-like mode of diapedesis but instead employ distinct mechanisms that may require progressive remodelling or disruption of the endothelial barrier.

**Fig. 1.**
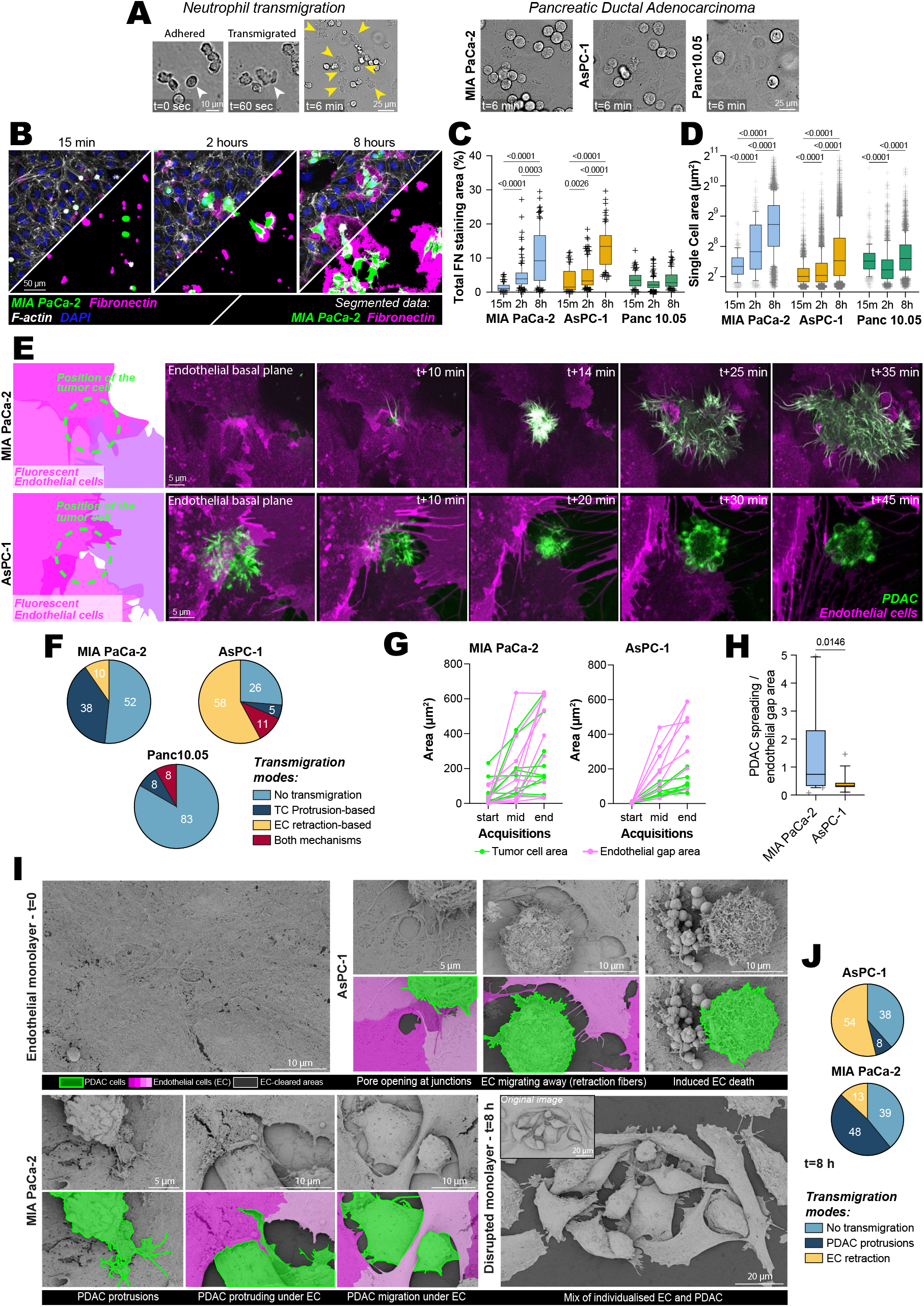
PDAC cells can breach the endothelial barrier via distinct modes of transmigration. (**A**) Brightfield images of neutrophils and PDAC cells, as indicated, perfused on top of an HUVEC endothelial monolayer. Still images of transmigrated neutrophils at 60 seconds and adhered cells at 6 minutes show numerous transmigrated neutrophils, whereas PDAC cells had not transmigrated within this time frame. Scale bars: 10 µm and 25 µm. (**B**-**D**) PDAC cells expressing Lifeact-mNeonGreen were added on top of endothelial monolayers and fixed after 15 minutes, 2 h, or 8 h. Samples were stained without permeabilisation with DAPI, phalloidin, and an anti-fibronectin antibody, and imaged on a spinning-disk confocal microscope. Cells and fibronectin-covered areas were then segmented and quantified. (**B**) Representative images of MIA PaCa-2 Lifeact-mNeonGreen cells and the corresponding cell and fibronectin labels are shown. Scale bar: 50 µm. (**C**) Quantification of accessible fibronectin (percentage coverage per field of view) for each cell line and time point (n > 72 fields of view; 3 biological replicates). (**D**) Quantification of single-cell area for each cell line and time point (n > 169 cells from 3 biological replicates). (**C, D**) Results are presented as boxplots (on a log2 scale), with whiskers extending from the 10th to the 90th percentiles. The boxes show the interquartile range, with the median indicated by a line. Data points outside the whiskers are shown as individual crosses. P values were determined using a Kruskal-Wallis test followed by Dunn’s post-test and are shown only for each cell line over time. (**E–H**) PDAC cells expressing Lifeact-mScarlet-I were added on top of a HUVEC endothelial monolayer expressing CAAX-EGFP (mosaic expression) and imaged using an Airyscan confocal microscope. (**E**) Representative still images extracted from live microscopy movies of MIA PaCa-2 and AsPC-1 transmigration events. Scale bar: 5 µm. (**F**) Pie charts quantifying the mode of transmigration per cell line. n = 31 cells for MIA PaCa-2, n = 19 cells for AsPC-1, and n = 36 cells for Panc 10.05. (**G**) Complete recordings of transmigration events were selected for further analysis of the relationship between cancer cell spreading and endothelial gap opening (n = 10 events for MIA PaCa-2 and n = 9 for AsPC-1). The cell area and the gap were manually measured at the start, middle, and end of each transmigration event (see Methods). (**H**) Using recordings of transmigration events, we calculated the ratio of PDAC expansion to gap area at the mid- and end-time points. A score close to 1, observed for MIA PaCa-2, indicates that the two dimensions are similar, whereas a score close to 0.5 for AsPC-1 indicates greater gap formation (n = 20 for MIA PaCa-2 and n = 18 for AsPC-1). P values were determined using a Mann-Whitney test. (**I**) MIA PaCa-2 and AsPC-1 cells were added on top of a HUVEC monolayer, fixed, and imaged using scanning electron microscopy. Representative images and corresponding segmentations of cancer and endothelial cells are shown. Scale bars as indicated; see panels. (**J**) Pie charts quantifying the mode of transmigration per cell line, based on SEM images. n (cells) = 23 for MIA PaCa-2 and n (cells) = 26 for AsPC-1. The raw numerical values and images used to make this figure have been archived on Zenodo.

To study this process, we selected three PDAC cell lines with distinct endothelial interaction phenotypes. Under flow, MIA PaCa-2 cells adhered efficiently across all tested flow speeds, AsPC-1 cells attached preferentially at lower flow rates but remained stably attached once adhered, and Panc 10.05 cells showed low endothelial engagement (12). We plated these three cell lines directly onto endothelial monolayers, thereby bypassing differences in initial arrest efficiency and enabling comparison of their post-attachment effects over the slower timescale of extravasation.

First, we analysed monolayer integrity by staining endothelial monolayers for fibronectin and thrombospondin without permeabilisation, to avoid visualising intracellular proteins and inaccessible basal ECM (Fig. 1B and Fig. 1C; Fig. S1A-C) (12, 25). Simultaneously, we measured cancer cell shape as a proxy for adhesion to the basal ECM (Fig. 1D; Fig. S1D). MIA PaCa-2 and AsPC-1 induced a significant, progressive exposure of the basal ECM, whereas Panc 10.05 had no effect (Fig. 1C; Fig. S1B). In addition, MIA PaCa-2 cells spread on the basal ECM within 2 hours and developed protrusive, elongated morphologies (reduced solidity), whereas AsPC-1 and Panc 10.05 cells remained rounded (higher solidity, smaller area; MIA PaCa-2 and AsPC-1 area, Fig. 1D; solidity of all lines, Fig. S1D).

Next, real-time impedance measurements of the endothelium during transmigration showed a progressive decrease in the presence of MIA PaCa-2 and AsPC-1, consistent with endothelial barrier disruption, whereas Panc 10.05 cells had minimal impact (Fig. S1E-F). Importantly, these cancer cell extravasation phenotypes observed in HUVEC monolayers were recapitulated in human pulmonary and dermal microvascular endothelial monolayers (HPMECs and HDMECs) (Fig. S1G-J), demonstrating that PDAC cell-line-specific phenotypes were conserved across endothelial models *in vitro*.

Taken together, PDAC transmigration occurs on significantly slower timescales than leukocyte diapedesis, varies substantially across PDAC lines, and markedly disrupts endothelial barriers.

### PDAC cells can transmigrate through multiple mechanisms

Next, we imaged PDAC transmigration at high resolution by plating Lifeact-mScarlet-I PDAC cells on a monolayer of CAAX-EGFP-expressing HUVECs (Fig. 1E and Fig. 1F; Supplementary Movie 1). The membrane-targeted CAAXEGFP signal delineated endothelial cell boundaries, allowing us to distinguish junctional opening from endothelial retraction. Consistent with our fixed-imaging results (Fig. 1C), most Panc 10.05 cells did not transmigrate and were unable to exploit the transient gaps that formed in the endothelial monolayer near arrested cells (Fig. S1K). In contrast, MIA PaCa-2 and AsPC-1 cells transmigrated efficiently, albeit by different mechanisms. MIA PaCa-2 cells extended thin, filopodia-like actin protrusions toward endothelial junctions to identify gaps, then rapidly spread on the underlying ECM (∼80% of events; Fig. 1F). These protrusions subsequently expanded, apparently widening the junctional opening into a gap large enough to accommodate the nucleus, usually within 35 minutes of initiation. This process coincided with pronounced endothelial deformation, including partial lifting of endothelial cells and the formation of contractile rings around the transmigrating cells (Fig. 1E and Fig. S1K). By contrast, AsPC-1 cells showed abundant short filopodia and blebbing at the endothelial surface but did not force their way through the junctions.

Instead, the underlying endothelial cells migrated away from the cancer cells, generating retraction fibres and locally detaching from the ECM, thereby creating space for AsPC-1 cells to access and spread on the basal ECM (∼90% of events, Fig. 1E and Fig. 1F; Fig. S1K). These behaviours occurred at both bi- and tri-cellular junctions, and transcellular transmigration was never observed.

These distinct extravasation modes were also reflected in the disparity between endothelial gap formation and cancer cell spreading over time (Fig. 1G-H). MIA PaCa-2 cell spreading and endothelial gap opening were closely linked. In contrast, AsPC-1 cells induced disproportionately larger gaps in the endothelial monolayer relative to their own spreading, consistent with an endothelial-retraction-driven opening mode. Notably, using an endothelial gap area equal to the initial projected area of the rounded cancer cell as the operational endpoint, both transmigration modes reached this endpoint on similar timescales after initiation (∼35–40 minutes; Fig. S1L), indicating that the mode of transmigration primarily influences the mechanism of barrier opening rather than its duration.

Finally, scanning electron microscopy further illustrated both transmigration modes (Fig. 1I and Fig. 1J), and classification of transmigration modes from randomly sampled SEM fields of view confirmed the distributions observed in live imaging (Fig. 1J).

Together, our data identify two PDAC transmigration modes: protrusion-led junctional invasion (MIA PaCa-2) and endothelial retraction or detachment (AsPC-1), both of which disrupt the endothelial barrier.

### PDAC transmigration is maintained on compliant and 3D matrices

Building on our observations on rigid glass, we imaged the transmigration process on hydrogels, which better recapitulate the physiological subendothelial mechanics. To assess the effect of basal substrate stiffness, we plated MIA PaCa-2 cells onto HUVEC monolayers grown on 10 kPa polyacrylamide gels and fixed the samples at 15 min, 2 hours, and 8 hours. Compared with glass, compliant substrates modestly reduced basal ECM accessibility during MIA PaCa-2 spreading, consistent with increased resistance of the endothelial monolayer to transmigration (Fig. S2A-B). Thus, reduced substrate stiffness attenuates PDAC-induced barrier disruption but does not abolish transmigration.

We next tested whether MIA PaCa-2 transmigration was associated with detectable mechanical deformation of the substrate. To address this, we combined live imaging with 2D traction force microscopy on bead-embedded polyacrylamide gels (see Methods). However, endothelial-generated lateral forces dominated the displacement field, making it difficult to resolve cancer cell-specific forces (Fig. S2C). Within these technical limitations, regions containing transmigrating MIA PaCa-2 cells generated lateral forces comparable to, or lower than, those measured in endothelial cells alone (Fig. S2D). Together, these data indicate that MIA PaCa-2 transmigration on compliant substrates is not associated with a clearly detectable increase in lateral traction forces.

Finally, we asked whether providing a three-dimensional basal matrix would enable endothelial resealing after PDAC transmigration. To test this, we grew HUVEC monolayers on collagen-fibronectin gels and recorded MIA PaCa-2 transmigration events using lattice light-sheet microscopy (Fig. S2E; Supplementary Movie 2). Despite the availability of basal space, we did not observe rapid closure of the endothelial monolayer, in contrast to previously described barrier-preserving endothelial remodelling programmes (9). To visualise matrix displacement during transmigration, we used CNA35-labelled collagen and analysed the 3D deformations (26). As in the 2D traction-force experiments, lateral displacements could not be attributed solely to cancer cells. However, axial displacement maps consistently revealed downward deformation of the matrix beneath transmigrating MIA PaCa-2 cells after they breached the endothelial barrier, indicating active invasion into the collagen matrix shortly after crossing the endothelium (Fig. S2E).

Together, these experiments show that PDAC-induced endothelial barrier disruption persists on compliant, three-dimensional matrices. Although substrate compliance modestly reduces ECM exposure, it does not prevent transmigration. Three-dimensional imaging reveals active downward invasion, with no evidence of rapid endothelial resealing during the first few hours of transmigration.

### Distinct modes of PDAC extravasation are recapitulated in zebrafish larvae

To determine whether the distinct extravasation modes observed *in vitro* are relevant *in vivo*, we imaged PDAC cell behaviour within the microvasculature of zebrafish larvae (27). We injected MIA PaCa-2, AsPC-1, and Panc 10.05 cells into the duct of Cuvier and assessed intravascular localisation and extravasation at 3 and 24 hours post-injection (Fig. 2A).

**Fig. 2.**
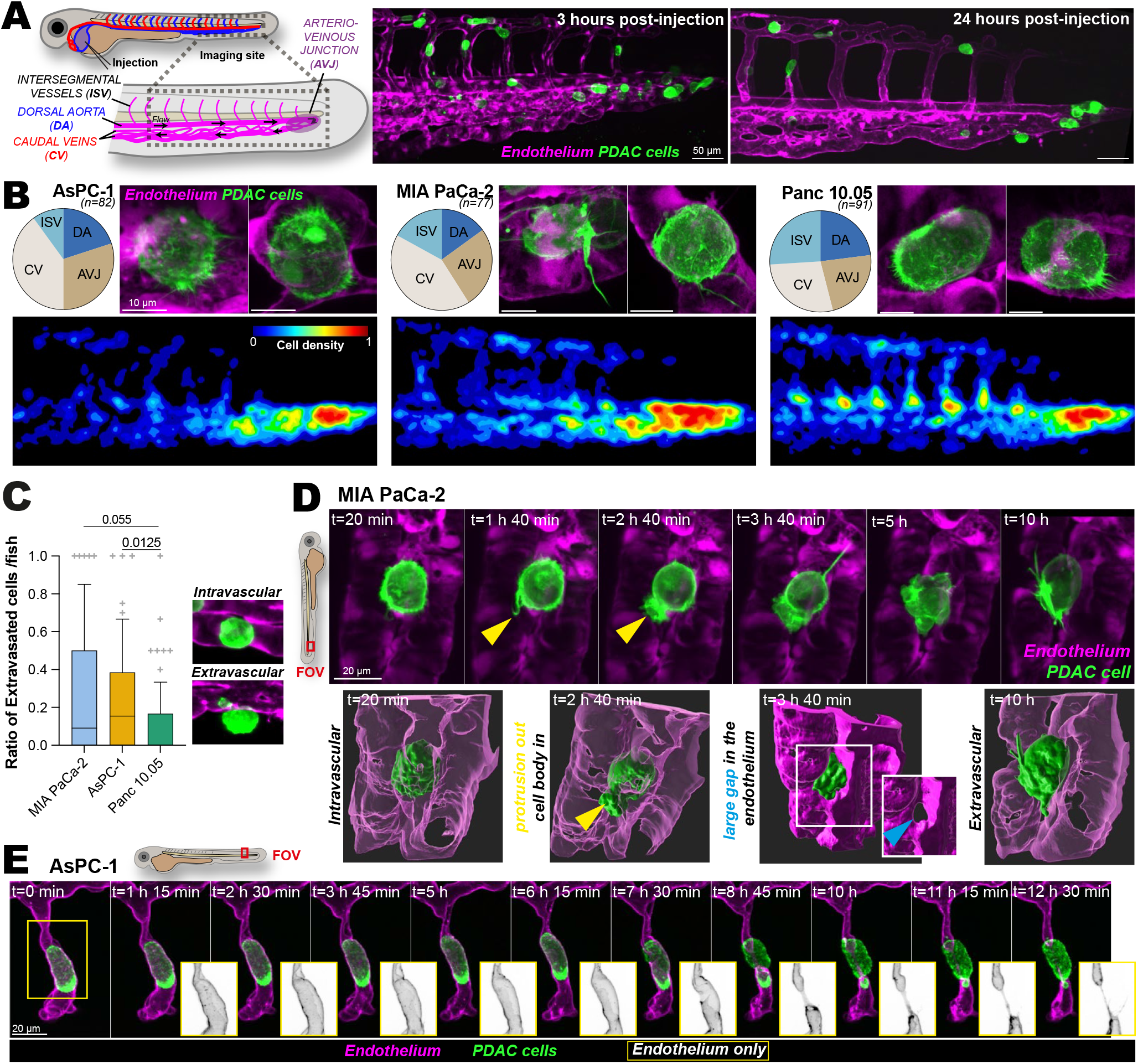
PDAC modes of extravasation are similar in vitro and in zebrafish larvae. (**A**) Schematic representation of PDAC cell injection into the duct of Cuvier of 48 hpf zebrafish larvae. Representative images show AsPC-1 Lifeact-mScarlet-I-expressing cells in the caudal plexus of *Tg(fli1:EGFP)* embryos at 3 h and 24 h post-injection. Images were acquired using a spinning disk confocal microscope. Scale bar: 50 µm. (**B**) Analysis of PDAC cell localisation and morphology in the caudal plexus at 3 h post-injection. Pie charts show the proportion of cells arrested in different anatomical regions for each PDAC cell line: ISV, intersegmental vessels; DA, dorsal aorta; AVJ, arteriovenous junction; CV, caudal veins. AsPC-1, MIA PaCa-2, and Panc 10.05: n = 82, 77, and 91 larvae, respectively, from 4, 3, and 4 biological replicates. High-magnification spinning-disk and Airyscan images show representative PDAC cell morphologies within the zebrafish vasculature. Scale bar: 10 µm. Arrest-site maps were generated by registering zebrafish larvae using DRMIME and segmenting arrested cancer cells using StarDist. Cancer-cell segmentation masks from all larvae were summed to generate stereotypical arrest maps for each PDAC cell line, displayed as normalised arrest-density maps. See Methods for details. (**C**) Quantification of the proportion of extravasated PDAC cells at 24 h post-injection. Representative images show examples of intravascular and extravascular cells used for classification. AsPC-1, MIA PaCa-2, and Panc 10.05: n = 87, 55, and 79 larvae, respectively, from 4, 3, and 3 biological replicates. Data are shown as boxplots. Boxes indicate the interquartile range, centre lines indicate the median, whiskers extend from the 10th to the 90th percentiles, and points outside the whiskers are shown as individual crosses. P values were determined using a Kruskal-Wallis test followed by Dunn’s post-test. (**D**) Still images from an overnight time-lapse recording of MIA PaCa-2 extravasation imaged using a spinning disk confocal microscope. The zebrafish schematic indicates the orientation of the field of view. Selected time points from 12 h of imaging were processed in Imaris to generate 3D renderings of key steps: protrusion extension, endothelial gap opening, and complete extravasation. Scale bar: 20 µm. (**E**) Still images from an overnight time-lapse recording of AsPC-1 extravasation imaged using a spinning disk confocal microscope. The zebrafish schematic indicates the orientation of the field of view. Selected time points from 13 h of imaging are shown. The black-and-white inset shows the endothelial channel, highlighting endothelial cell retraction over time. Scale bar: 20 µm. The raw numerical values and images used to make this figure have been archived on Zenodo (30).

At 3 hours post-injection, all three PDAC cell lines had arrested in the vasculature and displayed morphologies consistent with our *in vitro* observations: AsPC-1 cells exhibited numerous short filopodia; MIA PaCa-2 cells extended fewer but markedly longer protrusions along the vessel walls; and Panc 10.05 cells appeared larger, with polarised filopodia. To map stereotypical arrest sites across larvae, we registered zebrafish images using DRMIME (28) and segmented arrested cancer cells using StarDist (29) to generate population-level arrest maps (Fig. 2B; see methods). Mapping cancer-cell localisation within the caudal plexus showed that AsPC-1 and MIA PaCa-2 predominantly arrested near the arterio-venous junction, a site previously reported to favour cancer-cell arrest (Fig. 2B) (9). In contrast, Panc 10.05 cells were frequently detected at intersegmental vessel entry points, consistent with a greater propensity to remain trapped within narrower vessel segments (Fig. 2B) (16).

Across all PDAC lines, the total number of cancer cells per larva decreased by approximately 3- to 5-fold between 3 and 24 hours post-injection (Fig. 2A), indicating cell loss through recirculation and/or cell death. Among the remaining cells, endothelial crossing was inefficient, with average extravasation rates of around 20% for AsPC-1 and MIA PaCa-2 cells and close to 0% for Panc 10.05 cells (Fig. 2C). MIA PaCa-2 cells extended long protrusions through the vascular wall that grew into gaps, enabling extravasation (Fig. 2D; Supplementary Movie 3), whereas AsPC-1 cells extravasated concurrently with retraction of nearby endothelial cells (Fig. 2E; Supplementary Movie 3).

### AsPC-1 extravasation in the lung is associated with endothelial retraction

To define PDAC intravascular outcomes *in vivo* within a clinically relevant capillary bed, we injected EGFP- and luciferase-expressing MIA PaCa-2, AsPC-1, and Panc 10.05 cells into the tail vein of mice and monitored lung retention over time by combining bioluminescence imaging (Fig. 3A-C) with immunofluorescence imaging of whole-lung slices (Fig. 3A, D-E). Both approaches yielded concordant results across the three cell lines. AsPC-1 cells were retained in the lungs, with an initial decrease in bioluminescence followed by a lag phase at 1 day post-injection (Fig. 3B-C). By day 7, metastatic foci comprising tens of cells were observed (Fig. S3A), and by day 21, the bioluminescence signal increased exponentially, consistent with metastatic outgrowth (Fig. 3C; Fig. S3D). In contrast, MIA PaCa-2 bioluminescence was lost within 6 hours post-injection, and only limited tissue retention was detected by low-magnification confocal imaging (Fig. S3B, E). At 2 hours post-injection, these cells were apoptotic, as indicated by cleaved caspase-3 staining (Fig. S3B). Panc 10.05 bioluminescence declined more gradually and became undetectable by 2 days post-injection, consistent with the absence of fluorescent cells in lung sections (Fig. S3F). Unlike MIA PaCa-2 cells, Panc 10.05 cells were not prominently apoptotic but appeared fragmented within the tissue (Fig. S3C). We then performed systematic high-magnification imaging of the same lung slices to acquire 3D volumes and classify tumour cells as intravascular or extravasating/extravasated at 15 minutes, 6 hours, 24 hours, 48 hours, and 7 days post-injection (Fig. S3G-H). This analysis showed that AsPC-1 cells were the only cell line that consistently survived in circulation and extravasated from the lung vasculature into the tissue. A small number of MIA PaCa-2 and Panc 10.05 cells appeared to initiate extravasation in 6-hour post-injection samples, exhibiting protrusive morphologies and/or local endothelial perturbation, but these events did not result in sustained lung retention at later time points. This hierarchy was broadly consistent with MetMap, in which AsPC-1 showed strong lung colonisation, MIA PaCa-2 showed weak and contextdependent colonisation, and Panc 10.05 showed little or no lung colonisation (31).

**Fig. 3.**
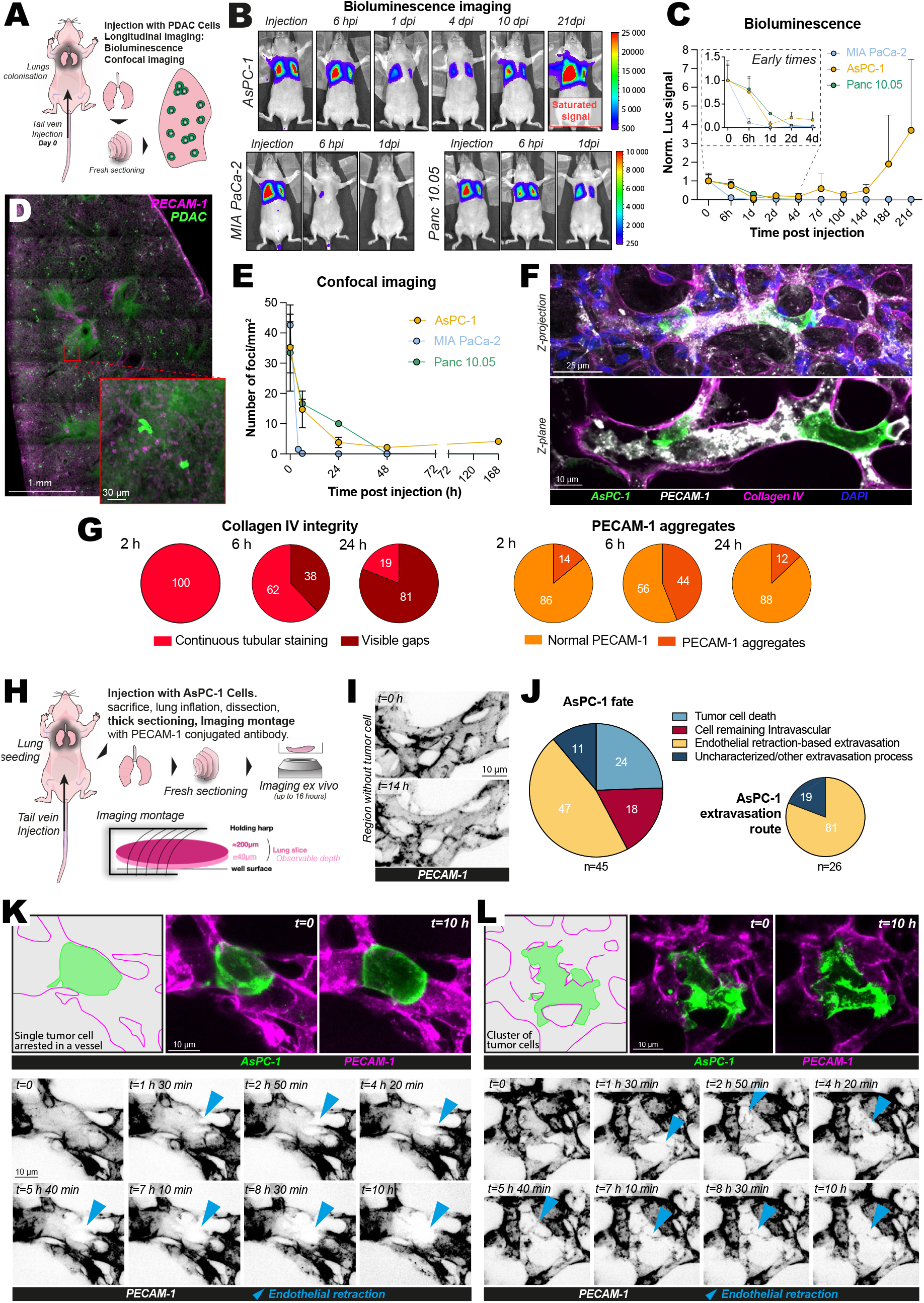
AsPC-1 cells induce local endothelial retraction during lung extravasation. (**A**-**G**) Luciferase- and eGFP-expressing MIA PaCa-2, AsPC-1, and Panc 10.05 cells were injected into nude mice via the tail vein. Lung retention and metastatic outgrowth were monitored by longitudinal bioluminescence imaging and endpoint fluorescence analysis of lung tissue. Initial animal numbers were as follows: AsPC-1, n = 17 from 3 independent experiments; MIA PaCa-2, n = 11 from 2 independent experiments; and Panc 10.05, n = 9 from 2 independent experiments. Animals were progressively sacrificed for tissue analysis; n values per time point are detailed in the Methods. (**A**) Schematic representation of the experiment. (**B**) Bioluminescence images at selected time points after injection for each PDAC cell line. A complete time course is shown for AsPC-1, whereas 0, 6, and 24 h time points are shown for MIA PaCa-2 and Panc 10.05 because the signal was no longer detectable after 24 h. Colour scales show raw bioluminescence counts. (**C**) Quantification of lung bioluminescence over time, normalised to the 15 min post-injection signal for each animal. Early time points from the same experiments are enlarged in the inset. (**D**) Representative low-magnification overview of a thick lung slice immunostained for PECAM-1. Tumour cells are fluorescently labelled. Scale bars: 1 mm in the overview and 30 µm in the inset. (**E**) Quantification of tumour-cell foci per mm^2^ of lung tissue over time, based on the fluorescent signal in lung sections. These data complement the bioluminescence analysis and show the number of tumour-cell foci retained in lung tissue over time. (**F**) High-magnification confocal z-projection and selected z-plane of AsPC-1 cells in the lung vasculature. Immunostaining for PECAM-1 and Collagen IV shows that AsPC-1 cells are lodged within the lung vasculature and that the endothelial layer is locally disrupted. PECAM-1 signal appears fragmented and detached from the Collagen IV-positive basal lamina. Scale bars: 25 µm and 10 µm (as indicated). (**G**) Quantification of Collagen IV gaps and PECAM-1 aggregation around AsPC-1 tumour cells using high-resolution z-stacks from the same lung slices as in (**D**). Number of foci analysed, from left to right: n = 81, 212, 21, 139, 277, and 16. **(H–L)** Lifeact-mNeonGreen-expressing AsPC-1 cells were injected into nude mice via the tail vein. Mice were sacrificed after tumour-cell injection, and lungs were extracted, thick-sectioned, labelled for PECAM-1, and imaged *ex vivo*. (**H**) Schematic representation of the experiment. (**I**) Representative images of *ex vivo* lung tissue at the start of imaging and after 14 h, acquired away from tumour cells. PECAM-1-conjugated antibody labelling indicates preservation of vascular structure during the imaging period. (**J**) Quantification of AsPC-1 cell behaviour in the lung vasculature during live *ex vivo* imaging. The first pie chart shows the distribution of all recorded foci. The second pie chart shows the prevalence of endothelial retraction among extravasation events. n = 45 foci from 8 biological replicates; n = 26 foci for the second pie chart. (**K, L**) Live imaging of AsPC-1 extravasation events in *ex vivo* lung slices. (**K**) Representative single-cell extravasation event. (**L**) Representative extravasation event involving a small AsPC-1 cluster. For each event, the schematic shows the starting configuration, merged images show the start and end points, and the time series of the PECAM-1 channel highlights local endothelial retraction around AsPC-1 cells. Scale bars: 10 µm. The raw numerical values and images used to make this figure have been archived on Zenodo.

We therefore focused on AsPC-1 cells to dissect their extravasation mechanism in mice. Lung sections collected at 2, 6, and 24 hours post-injection were co-stained for PECAM-1, which labels endothelial cells, and Collagen IV, which labels the basal lamina. Around arrested AsPC-1 cells, PECAM-1 staining revealed local endothelial detachment from the basal lamina and fragmentation into aggregated puncta (Fig. 3F; Supplementary Movie 4). We quantified the presence of Collagen IV gaps and PECAM-1 aggregation around AsPC-1 cells (Fig. 3G). Collagen IV gaps increased over time, whereas PECAM-1 aggregation peaked at 6 hours and resolved by 24 hours. These data suggest that AsPC-1 extravasation in the lung is associated with transient endothelial disruption and retraction. The same analysis of lungs from mice injected with MIA PaCa-2 cells did not reveal comparable vascular disruption (Fig. S3I).

To capture AsPC-1 extravasation dynamics directly in mammalian lung tissue, we performed *ex vivo* live imaging of precision-cut lung slices. Mice were injected with AsPC-1 cells, sacrificed approximately 1 hour post-injection, and their lungs were dissected, sectioned, labelled for PECAM-1, and imaged live immediately (Fig. 3H) (32, 33). Long time-lapse recordings were acquired from multiple fields of view per animal (Fig. 3I-L; Fig. S4). Importantly, PECAM-1 labelling showed that the vasculature remained intact away from tumour cells for at least 14 hours of imaging (Fig. 3I). Across the 45 recorded AsPC-1 foci, 24% of cells died within the vasculature, resembling the fragmentation phenotype observed for Panc 10.05 in fixed lung samples (Fig. 3J; Fig. S4A; Fig. S3C). 47% of cells extravasated via an endothelial-retraction-based mechanism (Fig. 3J-L; Fig. S4B; Supplementary Movies 5-7). The remaining cells were either intravascular (18%; Fig. S4C; Supplementary Movies 5-7) or extravasated through hybrid or difficult-to-classify mechanisms (11%). Notably, both single cells and small cell clusters induced local endothelial retraction (Fig. 3K-L; Fig. S4B). Finally, we prepared *ex vivo* lung slices at 24 hours post-injection and observed AsPC-1 cells migrating through the lung stroma, consistent with completion of extravasation (Fig. S4D; Supplementary Movie 8).

Together, these data show that AsPC-1 cells survived efficiently within the lung vasculature and could extravasate by inducing local endothelial retraction during the first few hours after injection. After breaching the endothelium, AsPC-1 cells entered the lung stroma within 24 hours, with a subset progressing to metastatic outgrowth by day 21. In contrast, MIA PaCa-2 and Panc 10.05 cells rarely completed productive extravasation and failed to sustain lung colonisation.

### AsPC-1 induces endothelial apoptosis during barrier failure

Our *in vitro*, zebrafish and *ex vivo* imaging consistently indicated that AsPC-1 transmigration/extravasation is associated with endothelial retraction and damage, prompting us to investigate whether endothelial cell death contributes to barrier failure.

We stained lung sections collected 6 hours post-injection for cleaved caspase-3 and quantified cleaved caspase-3-positive signals around or within tumour-cell foci in lungs injected with AsPC-1 or MIA PaCa-2 cells (Fig. 4A-B). Around AsPC-1 foci, 21% of cases showed nearby cleaved caspase-3 staining, whereas AsPC-1 cells themselves were rarely positive. By contrast, cleaved-caspase-3-positive neighbouring endothelial cells were rarely detected around MIA PaCa-2 foci (5%), whereas 77% of MIA PaCa-2 tumour cells were themselves cleaved-caspase-3-positive (Fig. 4B). These data suggest that AsPC-1 extravasation is associated with local apoptosis of neighbouring endothelial cells, consistent with endothelial stress or injury during barrier failure.

**Fig. 4.**
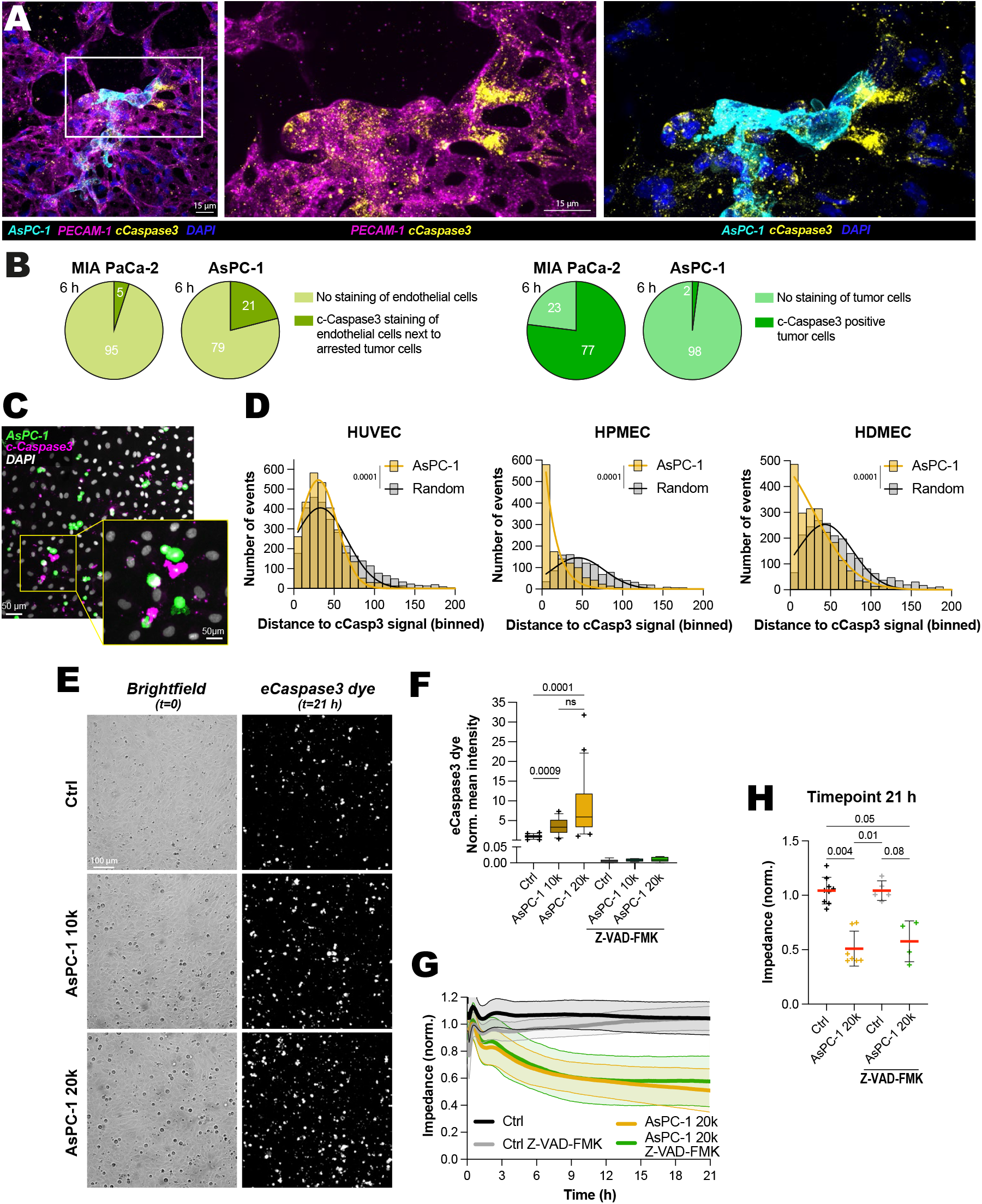
AsPC-1-induced endothelial apoptosis is dispensable for barrier disruption. (**A-B**) Nude mice were injected with AsPC-1 or MIA PaCa-2 cells, as in Fig. 3A, and their lungs were collected 6 h post-injection, fixed, sectioned, stained for cleaved caspase-3, and imaged by confocal microscopy. (A) Representative high-resolution confocal z-projection of extravasating AsPC-1 cells in mouse lung tissue. Scale bar: 15 µm. (**B**) Quantification of cleaved caspase-3 staining around tumour-cell foci and within PDAC cells. AsPC-1 foci were associated with a nearby cleaved-caspase-3-positive signal in 21% of cases, whereas MIA PaCa-2 cells were frequently cleaved-caspase-3-positive themselves. n = 48 foci for AsPC-1 and n = 22 foci for MIA PaCa-2. (**C-D**) AsPC-1 cells were added to confluent HUVEC, HPMEC, and HDMEC monolayers for 8 h, fixed, stained for cleaved caspase-3, and imaged by confocal microscopy. (**C**) Representative z-projection of an HPMEC monolayer after AsPC-1 addition and cleaved-caspase-3 staining. Scale bar: 50 µm. (**D**) Tumour cells and cleaved-caspase-3-positive endothelial cells were segmented, and nearest-neighbour distances were measured from each AsPC-1 cell to the closest cleaved-caspase-3-positive endothelial cell. Random point distributions generated in the same fields of view were analysed in parallel. Nearest-neighbour distance distributions are displayed as frequency plots. All values for each condition were compared using a Mann-Whitney test. n > 1,175 nearest-neighbour distances from 3 biological replicates. (**E-H**) 10,000 (10K) or 20,000 (20K) AsPC-1 cells were added to confluent HUVEC monolayers in xCELLigence plates in the presence of live fluorescent reporters for caspase-3 cleavage. Impedance and fluorescence were recorded in parallel for 21 h after addition of AsPC-1. Where indicated, endothelial monolayers were treated with Z-VAD-FMK (50 µM) to block caspase activity (n > 4 biological replicates). (**E**) Representative brightfield images at t = 0. Representative images of the caspase-3 reporter signal at 21 h after AsPC-1 addition. Scale bar: 100 µm. (**F**) Quantification of the caspase-3 reporter signal at 21 h after AsPC-1 addition. Data are shown as boxplots. Boxes indicate the interquartile range, centre lines indicate the median, and points outside the whiskers are shown as individual crosses. (**G**) Impedance traces recorded over 21 h after addition of AsPC-1 cells onto confluent HUVEC monolayers. (**H**) Endpoint impedance values at t = 21 h, plotted as mean ± SD with individual data points. (**F, H**) P values were determined using a Kruskal-Wallis test followed by Dunn’s post-test. The raw numerical values and images used to make this figure have been archived on Zenodo (30).

We hypothesised that AsPC-1-associated endothelial apoptosis could facilitate extravasation by creating space for tumour cells. To test this, AsPC-1 cells were added to HUVEC, HPMEC, or HDMEC monolayers, and cleaved caspase-3 immunostaining was used to identify apoptotic non-tumour cells after 8 hours (Fig. 4C). Distance analysis showed that apoptotic endothelial cells were significantly closer to AsPC-1 cells than expected from random spatial distributions, indicating that AsPC-1 cells locally promote endothelial apoptosis across endothelial models (Fig. 4D).

We next combined impedance measurements with live imaging of cell death reporters. AsPC-1 cells induced a cell number-dependent increase in caspase-3 reporter signal (Fig. 4E-F), consistent with apoptosis. To test whether apoptosis was required for barrier disruption, we added the pan-caspase inhibitor Z-VAD-FMK, which blocks caspase-mediated apoptosis (34). Barrier disruption was not diminished in the presence of Z-VAD-FMK, although the inhibitor blocked the AsPC-1-induced caspase-3 reporter signal (Fig. 4G-H). Thus, apoptosis accompanies AsPC-1-induced barrier disruption but is not required for loss of endothelial barrier integrity.

### AsPC-1 secreted factors induce endothelial barrier destabilisation

We next asked whether direct contact between AsPC-1 and endothelial cells is required to disrupt the endothelial barrier. To separate tumour cells from the endothelial monolayer while still allowing exchange of soluble factors, we co-cultured PDAC cells on Transwell inserts above endothelial monolayers and scored basal fibronectin exposure after 24 hours (Fig. 5A). Only AsPC-1 cells induced a strong increase in accessible fibronectin (Fig. 5B-C).

**Fig. 5.**
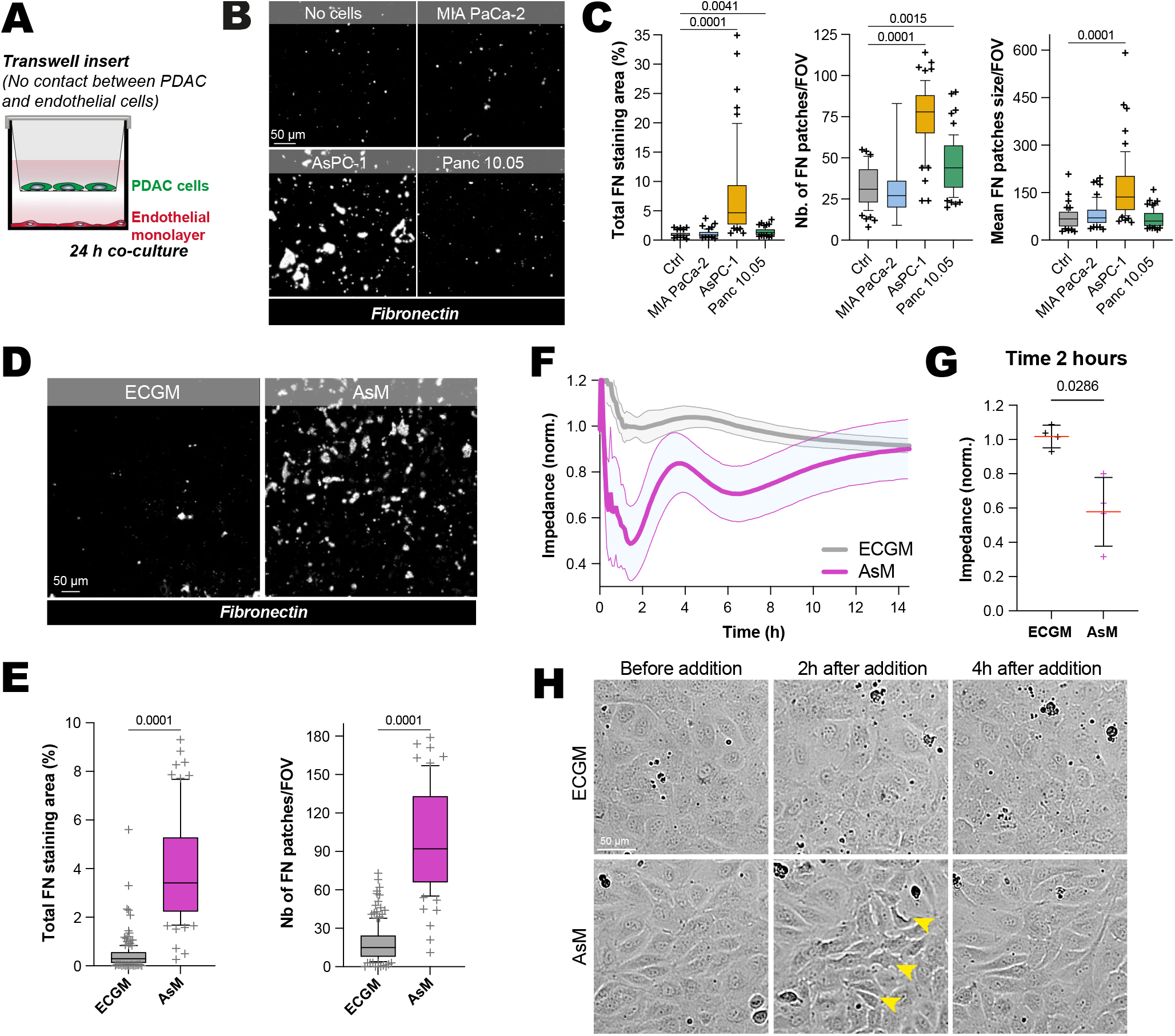
AsPC-1-secreted factors are sufficient to destabilise endothelial barriers. (**A-C**) PDAC cells were cultured on Transwell inserts (3 µm pores) above endothelial monolayers for 24 h, allowing exchange of soluble factors while preventing direct tumour-endothelial contact. Endothelial monolayers were then fixed and stained without permeabilisation with DAPI, phalloidin, and an anti-fibronectin antibody, and imaged by spinning-disk confocal microscopy. Fibronectin-positive areas were segmented and quantified. (**A**) Schematic representation of the Transwell experiment. (**B**) Representative images of accessible fibronectin staining. Only the segmented masks are shown. Scale bar: 50 µm. (**C**) Quantification of accessible fibronectin after 24 h, shown as percentage coverage per field of view, number of fibronectin patches, and mean patch size for each PDAC cell line (n > 101 fields of view; 6-8 biological replicates). (**D-E**) Concentrated conditioned media were prepared from AsPC-1 cells (AsM) and endothelial medium (ECGM) and applied to endothelial monolayers before fixation and fibronectin accessibility analysis. (**D**) Representative images of accessible fibronectin staining after 2 h treatment with concentrated media extracts diluted 1:10 in endothelial growth medium. Scale bar: 50 µm. (**E**) Quantification of the percentage coverage of accessible fibronectin and the number of fibronectin patches per field of view (n > 75 fields of view; 3-7 biological replicates). (**F-G**) Impedance measurements of confluent endothelial monolayers treated with concentrated media fractions. (**F**) Impedance traces were recorded for 14 h after addition of concentrated media. Data were normalised to the time of media addition (t = 0 h). n > 4 biological replicates. (**G**) Impedance values at t = 2 h, corresponding to the maximal decrease in impedance, plotted as mean ± SD with individual data points. P value was determined using a Mann-Whitney test. (**H**) Representative brightfield images of endothelial monolayers treated with concentrated medium or control medium, acquired immediately before media addition and at 2 h and 4 h after treatment. Scale bar: 50 µm. (**C, E**) Data are shown as boxplots. Boxes indicate the interquartile range, centre lines indicate the median, whiskers extend from the 10th to the 90th percentiles, and points outside the whiskers are shown as individual crosses. P values were determined using a Kruskal-Wallis test followed by Dunn’s post-test. The raw numerical values and images used to make this figure have been archived on Zenodo (30).

Next, we prepared conditioned medium from AsPC-1 cells grown to approximately 90% confluence, which were incubated for 24 hours in complete endothelial growth medium (AsM condition). Stimulation with AsPC-1 conditioned medium significantly increased fibronectin accessibility compared with endothelial growth medium alone (ECGM, Fig. 5D-E) within 2 h of stimulation. Real-time impedance measurements showed that this barrier perturbation occurred rapidly but was transient (Fig. 5F-G). Bright-field imaging of the same wells confirmed that the decrease in impedance coincided with visible disruption of the endothelial monolayer (Fig. 5H). AsM induced loss of endothelial cell-cell contacts and partial cell rounding (changes in image contrast, yellow arrows), and endothelial morphology largely recovered within 4 h of AsM addition, consistent with the rapid “pulse” effect observed in impedance measurements (Fig. 5F-H).

Together, these data show that direct contact between cancer cells and endothelial cells is not required for AsPC-1-induced endothelial barrier destabilisation. AsPC-1-secreted factors are sufficient to destabilise endothelial barrier integrity and transiently open endothelial monolayers.

### Saracatinib protects against AsPC-1-induced vascular disruption

Having identified endothelial barrier disruption as a key step exploited by AsPC-1 cells during extravasation, we next asked whether this vulnerability could be pharmacologically targeted. We used the rapid and most reproducible AsM-induced decrease in endothelial impedance to screen seven inhibitors targeting regulators of actomyosin tension, adhesion turnover, and junctional stability: non-muscle myosin II (blebbistatin), Src-family kinases (saracatinib), p38 MAPK

(SB202190), ROCK (H-1152), FAK (PF-573228), GSK3β (TWS-119), and mTORC1 (rapamycin). Endothelial monolayers were pretreated with the inhibitors for 2 hours, then challenged with AsM-conditioned medium in the continued presence of the inhibitor, and impedance was quantified 2 hours later, for a total inhibitor exposure of 4 hours (Fig. 6A-B; Fig. S5A). Strikingly, among the seven inhibitors tested, only saracatinib (35, 36) abolished the AsM-induced decrease in impedance without causing long-term disruption of the endothelial monolayer (Fig. 6B; Fig. S5A), pointing to saracatinib treatment as a strategy to protect against AsPC-1-induced barrier breakdown.

**Fig. 6.**
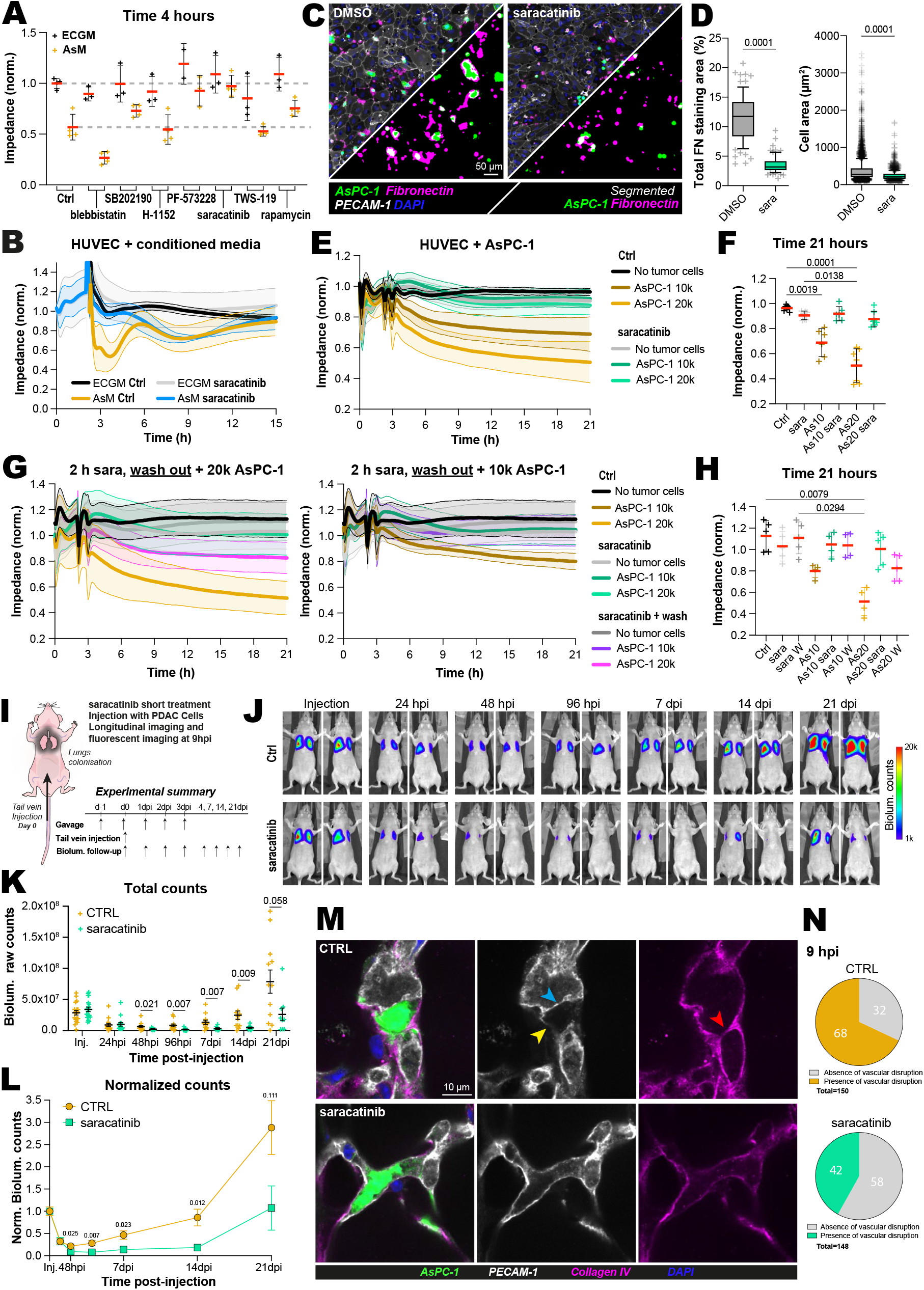
Saracatinib protects against AsPC-1-induced vascular disruption. (**A-B**) Endothelial monolayers were pre-treated for 2 h with inhibitors targeting cell adhesion, contractility, and cell-cell junction regulation, then challenged with AsM-conditioned medium in the continued presence of the inhibitors. Impedance was recorded over time and normalised to the time of inhibitor addition. (**A**) Impedance values at t = 4 h, corresponding to 2 h after AsM addition and the maximal impedance decrease, plotted as mean ± SD with individual data points. Complete impedance traces are shown in Fig. S5A and in (**B**) for saracatinib. n > 2 biological replicates. (**B**) Impedance traces were recorded over 15 h after addition of saracatinib to confluent endothelial monolayers. AsM-conditioned medium was added at t = 2 h. Data were normalised to the time of saracatinib addition (t = 0 h). n > 3 biological replicates. (**C-D**) AsPC-1 cells expressing Lifeact-mScarlet-I were added onto endothelial monolayers in the presence or absence of saracatinib and fixed after 8 h. Samples were stained without permeabilisation with DAPI, phalloidin, and an anti-fibronectin antibody, and imaged by spinning-disk confocal microscopy. AsPC-1 cells and fibronectin-positive areas were segmented and quantified. (**C**) Representative images and corresponding segmentation masks of AsPC-1 cells and accessible fibronectin. Scale bar: 50 µm. (**D**) Quantification of accessible fibronectin, expressed as percentage coverage per field of view, and AsPC-1 single-cell area in the presence or absence of saracatinib (n > 75 fields of view and n > 2222 cells; 3 biological replicates). (**E-F**) Confluent HUVEC monolayers were treated with saracatinib and challenged with AsPC-1 cells. Saracatinib was added at t = 0 h, and AsPC-1 cells were added 2 h later at the indicated densities. As10 and As20 correspond to 10,000 and 20,000 AsPC-1 cells per well, respectively. Sara indicates saracatinib treatment (n > 6 biological replicates). (**E**) Impedance traces recorded over 21 h after saracatinib addition. Data were normalised to the time of saracatinib addition (t = 0 h). (**F**) Endpoint impedance values at t = 21 h, plotted as mean ± SD with individual data points. (**G-H**) Confluent endothelial monolayers were pre-treated with saracatinib for 2 h. Saracatinib was then either maintained (sara) or washed out (sara W) before AsPC-1 cells were added at the indicated densities (n > 4 biological replicates). (**G**) Impedance traces recorded over 21 h after saracatinib addition, shown separately for each AsPC-1 density. Data were normalised to the time of saracatinib addition (t = 0 h). (**H**) Endpoint impedance values at t = 21 h, plotted as mean ± SD with individual data points. W indicates washout conditions. (**I-N**) Nude mice were treated with saracatinib or vehicle for 1 day before tail-vein injection of EGFP-luciferase-expressing AsPC-1 cells, and treatment was continued for 3 days after injection. Lung colonisation was monitored using longitudinal bioluminescence imaging, and early extravasation was assessed by endpoint analysis of lung tissue at 9 h post-injection. (**I**) Schematic representation of the *in vivo* saracatinib treatment and AsPC-1 injection protocol. (**J**) Representative bioluminescence images at selected time points after AsPC-1 injection. (**K-L**) Quantification of lung bioluminescence over time in vehicle- and saracatinib-treated mice. (**L**) Data were normalised to the early post-injection signal for each animal (n = 19 per condition at baseline; n = 12 in the control group and n = 9 in the saracatinib group at 3 weeks; see methods for details). (**M**) Representative high-resolution confocal images of AsPC-1 cells in lung tissue collected 9 h post-injection and immunostained for PECAM-1 and Collagen IV. Loss of PECAM-1 (yellow arrows), high-intensity PECAM-1 indicating the border of endothelial cells at the centre of the lumen (blue arrow), and collagen IV in the lumen (red arrow) are concordant signs of endothelial disruption. (**N**) Quantification of extravasation-associated vascular disruption around AsPC-1 cells based on PECAM-1 and Collagen IV integrity (n > 148 foci from 4 animals per condition). (**D**) Data are shown as boxplots. Boxes indicate the interquartile range, centre lines indicate the median, whiskers extend from the 10th to the 90th percentiles, and points outside the whiskers are shown as individual crosses. (**F, H, K, L**) Data are plotted as mean ± SD with individual data points shown where applicable. (**D, F, H, K, L**) P values were determined using a Kruskal-Wallis test followed by Dunn’s post-test. Raw numerical values and images used to generate this figure have been archived on Zenodo.

Focusing on saracatinib, we tested whether it could protect endothelial monolayers from AsPC-1-induced barrier disruption. Saracatinib markedly reduced fibronectin-positive endothelial gaps induced by AsPC-1 cells, and the AsPC-1 cells remained rounded on the monolayer surface, consistent with impaired transmigration (Fig. 6C-D; Fig. S5B). This protective effect was observed across endothelia of different origins: HUVEC, HPMEC, and HDMEC monolayers challenged with increasing numbers of AsPC-1 cells showed that saracatinib abolished the marked decrease in impedance seen under control conditions, indicating preserved endothelial barrier integrity and blocked transmigration (Fig. 6E-F; Fig. S5C-D). To determine whether saracatinib acts primarily on endothelial cells, on AsPC-1 cells, or on both, we pre-treated endothelial monolayers with saracatinib for 2 hours, washed out the inhibitor, and then added untreated AsPC-1 cells. Under these conditions, AsPC-1 cells induced only a moderate decrease in impedance, indicating that transient saracatinib exposure partially protects the monolayer. However, maximal barrier protection required continuous saracatinib exposure, suggesting that the full effect may involve sustained inhibition of endothelial responses and/or additional effects on tumour cells (Fig. 6G-H).

Given this robust *in vitro* protection, we next tested whether saracatinib could similarly limit AsPC-1 extravasation *in vivo* in the lung. Using established mouse-dosing protocols for saracatinib as a guide (37, 38), we designed a treatment schedule to target the early extravasation window (Fig. 6I). Mice received saracatinib the day before tail-vein injection of AsPC-1 cells, again 4 hours before injection on the day of injection (36, 39), and then once daily for 3 additional days after injection, for a total of 5 days of treatment. Longitudinal bioluminescence imaging showed that saracatinib delayed AsPC-1 lung outgrowth, with treated mice exhibiting delayed regrowth over 21 days compared with controls (Fig. 6J-L). To directly capture the early extravasation step, we collected lungs 9 hours post-injection and stained them for PECAM-1 and Collagen IV (Fig. 6M-N). Quantification of vascular integrity around AsPC-1 cells revealed a 38% reduction in extravasation-associated vascular disruption in saracatinib-treated mice (Fig. 6M-N).

Together, our *in vitro* and *in vivo* data show that Saracatinib protects endothelial barriers from AsPC-1-induced destabilisation, reduces early extravasation-associated vascular disruption in the lung, and delays subsequent metastatic outgrowth. These findings establish AsPC-1 extravasation as a pharmacologically targetable step in metastatic dissemination and support the use of endothelial barrier protection as a therapeutic strategy to limit metastatic colonisation.

## Discussion

Here, we show that PDAC extravasation is not uniform and can proceed through at least two distinct modes. MIA PaCa-2 cells use protrusions, extending actin-rich structures toward endothelial junctions or local openings before spreading beneath the monolayer. In contrast, AsPC-1 cells induce endothelial retraction, remaining rounded while adjacent endothelial cells withdraw, detach from the basal lamina, and open the barrier around the tumour cell. This heterogeneity may be therapeutically important, as it suggests that a single strategy targeting one extravasation mode is unlikely to be effective across different PDAC tumours.

Building on our previous work characterising endothelial remodelling around arrested cancer cells in breast cancer (9), our findings show that PDAC extravasation also diverges from the classical analogy of leukocyte diapedesis. Leukocytes traverse the endothelium through adhesion, crawling, paracellular or transcellular passage, and rapid barrier resealing (15, 40). By contrast, PDAC cells can remain apically associated with the endothelium for hours before barrier opening begins; once initiated, the operational gap-area endpoint was reached within approximately 35-40 min, accompanied by exposure of the basal ECM. These characteristics position PDAC at the barrier-disruptive end of the tumour-extravasation spectrum, where increased endothelial permeability, junctional destabilisation, and basement membrane perturbation accompany cancer cell escape (14, 15).

Our data further distinguish vascular arrest from productive extravasation. Panc 10.05 cells were effectively arrested within zebrafish vessels and lung capillaries, but rarely crossed the endothelium. Consistently, MetMap identified strong lung colonisation by AsPC-1, weak and context-dependent colonisation by MIA PaCa-2, and little or no colonisation by Panc 10.05 (31). Our data support intravital imaging studies indicating that arrest, survival, endothelial crossing, and metastatic outgrowth are distinct steps rather than an inevitable sequence (4, 7, 41). Identifying which adhesive, mechanical, or secretory properties enable an arrested PDAC cell to become transmigratory remains an important direction for future work. We favour this step-wise dissection of the metastatic cascade, as it can reveal therapeutic windows that endpoint metastatic-burden assays alone would miss.

MIA PaCa-2 cells exemplify a junction-probing mode of extravasation. Their filopodia-like, actin-rich protrusions probe endothelial junctions and local gaps before they spread onto the basal ECM. This behaviour aligns with previous evidence that transendothelial migration can rely on active adhesion and protrusive engagement with endothelial junctions and the subendothelial matrix, mediated by Cdc42-β1-integrindependent adhesion and invadopodia formation (14, 42, 43).

In this mode, cancer cells drive their own exit by identifying and enlarging a permissive opening in the endothelium. Understanding how these protrusions detect junctional weakness and engage the basal matrix will be central to elucidating protrusion-led extravasation.

AsPC-1 cells exhibit a distinct mode of extravasation in which tumour-endothelial signalling, rather than cancer-cell protrusions, predominates. Instead of traversing endothelial junctions, AsPC-1 cells remain rounded while neighbouring endothelial cells retract and detach. This behaviour is conceptually similar to endothelial-remodelling processes such as pocketing and angiopellosis-like events, in which endothelial cells reshape around arrested tumour cells or clusters (9, 15, 44). However, the AsPC-1 phenotype is not barrier-preserving: it involves endothelial detachment, cell death, and exposure of the basal ECM, thereby phenotypically resembling tumour-cell-induced endothelial necroptosis (21). Although AsPC-1 cells induce endothelial apoptosis, pan-caspase inhibition does not prevent barrier opening, indicating that cell death is dispensable for this process. Accordingly, AsPC-1-induced endothelial apoptosis appears to be a downstream consequence of retraction-associated stress rather than the initiating mechanism. Whether the resulting vascular microlesions cause blood leakage or recruit inflammatory cells and platelets remains unknown.

We report that tumour-derived soluble cues mediate endothelial retraction, at least in part. Conditioned medium and transwell experiments show that direct tumour-endothelial contact is not required for AsPC-1 cells to destabilise endothelial monolayers, indicating that secreted factors act upstream of barrier failure. This finding extends earlier work showing that conditioned media from PSN-1 cells increased endothelial permeability and enhanced pancreatic cancer-cell adhesion and invasion (45). The identity of the AsPC-1-derived factors remains unknown, but candidates include VEGF-family ligands, inflammatory mediators, proteases, and bioactive lipids and metabolites, and most likely several factors acting synergistically to affect endothelial junctions.

We identify saracatinib treatment as a potent means of preserving endothelial barrier integrity in response to AsPC-1 cells.

However, because saracatinib was administered systemically, its *in vivo* effects cannot be assigned specifically to endothelial cells and may also involve tumour cells or other host compartments. This finding aligns with the established roles of Src-family kinases in regulating endothelial permeability and junction remodelling. VEGF-driven Src signalling disrupts endothelial barriers and promotes tumour-cell extravasation and metastasis, while VEGFR2-dependent junctional signalling contributes to vascular permeability and metastatic spread (46, 47). Src and Pyk2 also regulate ICAM-1-dependent VE-cadherin phosphorylation during leukocyte transendothelial migration, positioning Src-family kinases as central regulators within broader endothelial junction-remodelling programmes (48).

The divergent behaviours of MIA PaCa-2 and AsPC-1 cells suggest that the mode of extravasation may be linked to tumour-cell state along the mesenchymal-amoeboid axis. MIA PaCa-2 cells exhibit a protrusive, mesenchymal-like phenotype, whereas AsPC-1 cells remain rounded and bleb-rich, resembling an amoeboid-like state characterised by low adhesion, cortical RhoA/C-ROCK-myosin II contractility, and bleb-based protrusions (49–52). Notably, amoeboid behaviour is not solely a mechanical state: cytokine signalling through GP130-JAK1-STAT3 can reinforce ROCK-myosin-II-dependent contractility, and rounded myosin-II-high PDAC cells have been linked to invasion, metastasis, and a CD73-PI3K-RhoA-ROCK-dependent immunomodulatory secretome (53, 54). The observation that AsPC-1-conditioned medium destabilises endothelial monolayers extends this concept to the vascular wall and suggests that rounded PDAC states may couple mechanical plasticity to paracrine remodelling of the endothelium.

Together, these findings show that PDAC cells breach endothelial barriers through mechanistically heterogeneous modes rather than a single leukocyte-like programme. This heterogeneity has direct therapeutic and potentially diagnostic implications. A protrusion-led mode, as used by MIA PaCa-2 cells, and a retraction-driven mode, as used by AsPC-1 cells, are unlikely to share a single vulnerability. Our identification of saracatinib as a mode-specific strategy against AsPC-1induced barrier disruption, both *in vitro* and *in vivo*, provides proof of concept that extravasation can be pharmacologically targeted. Building on earlier studies, the idea that amoeboid and mesenchymal cell states may distinguish extravasation modes can now be tested using retrospective archived clinical samples (55). Future work should determine how prevalent each extravasation mode is across PDAC patient populations, whether cell morphology more broadly predicts the extravasation mode, and whether protrusion-led extravasation is susceptible to a distinct, complementary therapeutic strategy. More broadly, our findings suggest that strategies protecting the endothelial barrier, tailored to the dominant mode of vascular breach, may offer a viable and currently underexploited avenue for limiting metastatic dissemination.

## Methods

### Cells

Pancreatic ductal adenocarcinoma (PDAC) cell lines AsPC-1 (ATCC-CRL-1682), MIA PaCa-2 (ATCC-CRL1420), and Panc 10.05 (ATCC-CRL-2547) were cultured under standard conditions in 10 cm plastic culture dishes. MIA PaCa-2 cells were maintained in DMEM high-glucose (DMEM HG), whereas AsPC-1 and Panc 10.05 cells were maintained in RPMI 1640. All media were supplemented with 10% fetal bovine serum (FBS) (BioWest, S1810), 1% L-glutamine (Sigma-Aldrich, G7513), and 1% penicillin-streptomycin (Sigma-Aldrich, P0781). Panc 10.05 medium was additionally supplemented with 10 U/mL human insulin (Sigma-Aldrich, I0908). PDAC cell lines were authenticated at the start of the project by the Leibniz Institute DSMZ (Deutsche Sammlung von Mikroorganismen und Zellkulturen) using STR profiling. HEK293FT cells (for lentiviral production) were cultured in DMEM HG supplemented with 10% FBS, 1% L-glutamine, and 1% penicillin-streptomycin. All cell lines were routinely tested for mycoplasma infection and were negative.

Human umbilical vein endothelial cells (HUVECs; PromoCell, C-12203) were cultured in ready-to-use Endothelial Cell Growth Medium (ECGM; PromoCell, C-22010 and C-39215) supplemented with 1% penicillin-streptomycin (Sigma-Aldrich, P0781). Human pulmonary microvascular endothelial cells (HPMECs; PromoCell, C-12281) and human dermal microvascular endothelial cells (HDMECs; PromoCell, C-12212, adult donors) were cultured in ready-to-use Endothelial Cell Growth Medium MV (ECGM MV; PromoCell, C-22020 and C-39225) supplemented with 1% penicillin-streptomycin (Sigma-Aldrich, P0781). The three primary endothelial cell types were obtained at P0 (commercial vial), expanded to P3, and stored at -150°C until use. Vials were thawed, cells were transferred to 10-centimetre culture dishes, and P4 cells were grown for at least 2 days to near confluence before being plated in appropriate vessels for each experiment (P5 cells were used in experiments). HUVECs expressing membrane-targeted EGFP (CAAX-EGFP) were generated by lentiviral transduction (see next section). Briefly, we thawed the P0 commercial vial and infected the cells with virus-containing medium the following day. Cells were expanded from P0 to P3, yielding stock vials containing 50% fluorescent cells, which were used to grow mosaic monolayers.

### Generation of fluorescent reporter cell lines

EGFP-Luc, lifeact-mNeonGreen, and lifeact-mScarlet-I-expressing AsPC-1, Panc 10.05, and MIA PaCa-2 cell lines were generated as previously described (12) using a third-generation lentiviral system. HEK293FT cells (Thermo Fisher Scientific, R70007) were used to produce lentiviral particles. Cells were transfected with a third-generation lentiviral packaging system comprising the envelope plasmid pMD2.G (a gift from Didier Trono, Addgene 12259), packaging plasmids pMDLg/pRRE (a gift from Didier Trono, Addgene 12251, (56)) and pRSV-Rev (a gift from Didier Trono, Addgene 12253, (56)), and the respective target sequence plasmids pLV430G-ofl-T2A-EGFP (57, 58), pLenti6.3/TO/V5-DEST-Lifeact-mNeonGreen (Addgene 225494), and pLenti6.3/TO/V5-DEST-Lifeact-mScarlet-I (Addgene 225495).

HUVECs expressing membrane-targeted EGFP (EGFP-CAAX) were generated by lentiviral transduction. The pLenti6.3/TO/V5-DEST-EGFP-CAAX was prepared by PCR-amplifying the EGFP-CAAX fragment from a pEGFP-C1-CAAX plasmid and adding flanking KpnI and XhoI restriction sites, using the following primers: EGFPcaax_F 3’- ATCCGGTACCGTCGCCACCATGGTGAGCAAGGGCG-5’ and EGFPcaax_R 3’- TGAGCTCGAGATCTGGATCCTCAGGAGAG-5’. After enzymatic digestion of the PCR product with KpnI (Thermo Fisher Scientific, FD0524) and XhoI (Thermo Fisher Scientific, FD0694), and of the pENTR2b entry vector (Thermo Fisher Scientific, A10463), followed by ligation, we obtained pENTR2b-EGFP-CAAX. An LR reaction (LR Clonase II; Invitrogen, 56485) was then carried out following the manufacturer’s instructions with the pLenti6.3/TO/V5-DEST destination vector (Thermo Fisher Scientific, V53306). We verified plasmids by sequencing and analytical digestion.

### Isolation of neutrophils from human blood samples

Human blood was collected from healthy adult volunteers in heparin-treated tubes with ethical permission from VARHA (ETMK Dnro: 43/1801/2015). Neutrophils were isolated from whole blood by density-gradient centrifugation using Percoll gradients (Sigma-Aldrich, p1644). Neutrophils and red blood cells were collected from beneath the Percoll layer. Red blood cells were removed by osmotic lysis: 1 mL of 0.2% NaCl was added to the cell pellet and vortexed for 18 seconds, followed by 1 mL of 1.8% NaCl. The cells were then pelleted. This step was repeated three times. Neutrophil fractions were washed with PBS, kept on ice, and used within 3 h of isolation.

### Antibodies and reagents

The following primary antibodies were used in this study for immunofluorescence (IF): anti-fibronectin (FN) (1:200, Sigma-Aldrich, F3648); anti-thrombospondin (1:200, Abcam, ab1823); anti-PECAM-1 (MEM-5, 1:200, Invitrogen, 37-0700); anti-PECAM-1 (Mec13.3, 1:100, BioLegend, 102501); Alexa Fluor-488- or -647-conjugated antibodies; anti-Collagen-IV (1:200, Abcam, ab19808); anti-cleaved caspase-3 (1:200, Cell Signaling Technology, 9664); and anti-cleaved caspase-3 (1:200, Cell Signaling Technology, 9661).

For immunofluorescence, secondary antibodies were used at a 1:400 dilution and were all purchased from Invitrogen. These included Alexa Fluor 647-conjugated anti-mouse IgG (A21235), Alexa Fluor 488-conjugated anti-mouse IgG (A11001), Alexa Fluor 568-conjugated anti-mouse IgG (A10037), Alexa Fluor 488-conjugated anti-rat IgG (A21208), Alexa Fluor 568-conjugated anti-rat IgG (A11077), Alexa Fluor 647-conjugated anti-rabbit IgG (A21244), Alexa Fluor 488-conjugated anti-rabbit IgG (A11008), and Alexa Fluor 568-conjugated anti-rabbit IgG (A10042).

Additional reagents used in this study included DAPI (1:5,000; Thermo Fisher Scientific, D1306), Alexa Fluor 647-conjugated phalloidin (1:800; Invitrogen, A30107), Pro-Long Glass mounting medium (Thermo Fisher Scientific, P36980), eCaspase-3 Blue reagent (2.5 µM in PBS; Agilent, 8711027), eAnnexin V Blue reagent (0.125 µg/mL in ECGM; Agilent, 8711026), CNA35-HaloTag, generated in-house (59), Janelia Fluor 646 HaloTag ligand (200 nM; Promega, HT1060), SPY-650-Fastact (1:2,000; Spirochrome, SC502), poly-D-lysine (10 µg/mL; Gibco, A3890401), fibronectin (10 µg/mL; Sigma-Aldrich, 341631), rat-tail collagen (Sigma-Aldrich, 08-115), IL-1β (10 ng/mL; RD Systems, 201-LB), blebbistatin (20 µM; STEMCELL Technologies, 72402), saracatinib (Selleckchem, S1006), used at 5 µM in ECGM *in vitro* and at 50 mg/kg/day by oral gavage in 0.5% carboxymethylcellulose *in vivo*, with the stock solution prepared in DMSO, TWS-119 (5 µM; Selleckchem, S1590), rapamycin (2 µM; Santa Cruz Biotechnology, sc-3504A), PF573228 (5 µM; Selleckchem, S2013), SB202190 (10 µM; Bio-Mediator/MedChemExpress, HY-10295), H-1152 (10 µM; Calbiochem/Merck Millipore, 555550), Z-VAD-FMK (50 µM; MedChemExpress, HY-16658B), puromycin (1 µg/mL; MedChemExpress, HY-B1743), horse serum (Gibco, 16050-122), bovine serum albumin (BSA; Sigma-Aldrich, 126575), Triton X-100 (Sigma-Aldrich, 1000882545), sodium azide (Sigma-Aldrich, 71289-5G), Dulbecco’s phosphate-buffered saline (DPBS; Biowest, L0615-500), and D-luciferin (150 mg/kg, administered intraperitoneally; MedChemExpress, HY-12591).

### Endothelial cell monolayer preparation

Microfluidic channel slides, 1-channel µ-Slide LUER 0.4 (Ibidi, 80176), Ibidi 8-well µ-Slide Luer Glass Bottom (Ibidi, 801779), 96-well plates with gold electrodes for impedance measurements, E-Plate VIEW 96 (Agilent, 300601020), E-Plate 96 (Agilent, 5232368001), and glass coverslips in 24-well cell culture plates were used in this study. Before seeding endothelial cells, vessels were coated with 10 µg/mL fibronectin (Sigma-Aldrich, 341631) for 1 hour at 37 °C. Endothelial cells were seeded at 5 × 10^5^ to 8 × 10^5^ cells/mL and cultured with media changes daily in wells or twice daily in channels for 3-4 days. A peristaltic pump (Masterflex® Ismatec® 78018-24) was used to create flow for the perfusion experiment. The flow system, including the tubing assembly, was custom-built in-house using components from Ibidi and IDEX: ISMATEC-070535-04i-ND SC0052T, IBIDI-10840, IBIDI-10842, IBIDI-10829, IBIDI-10802, and IBIDI-10827. Detailed assembly instructions are provided in Osmani et al. (60). The microfluidic experimental dataset (Fig. 1A) was generated previously (12). See the impedance section for additional details on E-Plate preparation.

### Hydrogel preparation, surface activation and Traction Force Microscopy

35 mm glass-bottom dishes (D35-14-1, Cellvis) were treated for 30 min at room temperature with 1 mL of bind-silane solution (7.14% Plus One Bind Silane (Sigma) in 7.14% acetic acid in absolute ethanol), followed by two washes with 2 mL of absolute ethanol and air-drying. To make the hydrogels (10 kPa), 1.7 µL of sonicated fluorescent beads (FluoSpheres™ Carboxylate-Modified Microspheres, 200 nm Yellow-Green, F8811) were added to 500 µL of hydrogel mix containing 94 µL of 40% acrylamide (Sigma) and 50 µL of N,N’-methylenebisacrylamide (M1533, Sigma) in PBS. 5µL of 10% ammonium persulfate (1610700, Bio-Rad) and 1µL of N,N,N’N,’-tetramethylethylenediamine (T9281, Sigma) were added to initiate polymerisation, and the mixture was rapidly vortexed. Then, 11.8 µL was added to the dishes, and a 13-mm glass coverslip was placed on top. Hydrogels were left to polymerise at room temperature for 1 h before PBS was added and the glass coverslips were removed. For surface activation, hydrogels were incubated with 500 µL of 0.2 mg/mL Sulfo-SANPAH (803332, Sigma) and 2 mg/mL N-(3-Dimethylaminopropyl)-N-ethylcarbodiimide hydrochloride (03450, Sigma) in 50mM HEPES for 30 min at room temperature with gentle agitation, followed by UV irradiation for 10 min. Hydrogels were washed four times with sterile PBS before coating with 10 µg/mL fibronectin.

Endothelial cells were seeded at 800,000 cells/mL and cultured for 3 days until confluence. Cells were labelled with SPY-650-Fastact (1:2000) for 4 h, washed with medium, and placed in the thermostated chamber of the Marianas spinning disk microscope (see the dedicated section for details; objectives used were ×40, NA 1.1, Water, Zeiss LD C-Apochromat). Then, 50,000 Lifeact-mScarlet-I-expressing MIA PaCa-2 cells were added, and multi-channel live imaging was performed for 2 hours. Cells were removed by adding 20 µL of 20% SDS, after which a final acquisition was performed for traction force microscopy. To correct for drift, bead and cell images acquired before and after cell removal were aligned using the Fast4Dreg plugin in Fiji (61). Bead tracking and force measurements were analysed using TFM software (62) in MATLAB (MathWorks, version 2024a). The displacement field was calculated on a 100 × 100 grid using the PIV suite, with subpixel correlation via image interpolation, a template size of 21 pixels, and a maximum pixel displacement of 20. For displacement-field correction, vector-field outliers were filtered, and a normalised displacement residual threshold of 2 was applied. Force-field calculation was performed using Fourier Transform Traction Cytometry with a regularisation parameter of 0.0001. MIA PaCa-2 cell masks were generated from the red-channel images using FIJI/ImageJ (63) and manually classified as corresponding to cells that underwent transmigration or did not transmigrate. The masks were overlaid onto traction maps in R (A Language and Environment for Statistical Computing. R Foundation for Statistical Computing, Vienna, Austria. https://www.R-project.org/). Mean pixel values inside and outside the masks were calculated and plotted.

### Measuring ECM displacement during transmigration

To measure the displacements exerted by HUVECs and PDAC cells on the underlying ECM in 3D, substrates consisting of type I collagen and fibronectin were prepared on glass-bottom dishes (Cellvis, D35-14-1-N). For each gel, 90 µL of rat tail collagen (Sigma-Aldrich, 08-115) in acetic acid was mixed with 10 µL of cold 10× PBS neutralised by adding 1 µL of 1 M NaOH, then combined 1:1 with a cold fibronectin solution (0.5 mg/mL in PBS) and spread evenly over the glass bottom. The gels were allowed to polymerise for 45 min at 37 °C in a humidified incubator, then immersed in PBS and stored at 4 °C. Transduced HUVECs expressing CAAX-EGFP (mosaic) were grown as a monolayer, as described above, for 3 to 4 days on the gels. CNA35-HaloTag and 646 Halo ligand were used to stain the collagen network for 4 hours before transfer of the sample to a thermostatted chamber of a lattice light-sheet microscope (Zeiss; see technical details in the dedicated section). MIA PaCa-2 cells expressing Lifeact-mScarlet-I were added, and imaging was started in selected regions showing a high density of CAAX-EGFP-expressing HUVECs and PDAC cells on top for 3-hour time-lapse imaging.

3D displacement fields were calculated using TFMLAB (64, 65), a traction force microscopy toolbox implemented in MATLAB R2022a (MathWorks). Collagen fibres and MIA PaCa-2 cells were segmented using the CNA35 and Lifeact signals, with variable threshold adjustment and a minimum object size of 5-10 × 10^3^ voxels for the cells. Rigid image registration was performed using a gradient-based algorithm. Displacements were measured from CNA35 images using a 10 × 10 × 10-pixel/voxel grid, the default registration metric and optimiser, and a second post-shift correction. Results were visualised using ParaView v5.11.0 (66). Lateral and axial displacements were also visualised as raster and quiver plots using R (v 4.2.2) and RStudio (v 2024.04.0).

### xCELLigence Impedance measurement

Impedance measurements were conducted using the xCELLigence Real-Time Cell Analyser DP Instrument (W380; ACEA). 96-well plates (E-Plate VIEW 96, Agilent, 300601020; E-Plate 96, Agilent, 5232368001) were coated with fibronectin (Sigma-Aldrich, 341631) at 10 µg/mL for 1 hour at 37 °C. After a PBS wash, each well was filled with endothelial cell growth medium (ECGM or ECGM MV), and an initial impedance reading was recorded to establish a baseline before adding endothelial cells. Impedance was then recorded every 30 minutes to 1 hour over 72 hours. The culture medium was refreshed every 24 hours to maintain optimal cell growth conditions. After 3 days, 10,000 or 20,000 PDAC cells per well were added to the confluent monolayers, and acquisition parameters were adjusted to 5- to 15-minute intervals for impedance measurements and 30-minute intervals for imaging (brightfield and fluorescence). Impedance was measured for an additional maximum of 24 hours. The same protocol was used for experiments with concentrated media (see below), in which 1:10 media extracts were pipetted into the well. For inhibitor treatments, inhibitors were added 2 hours before PDAC cells or media extract and were either maintained at constant concentrations or washed out with fresh media before the addition of AsPC-1 cells. For experiments involving the live dye eCaspase-3 Blue reagent in PBS (2.5 µM, Agilent, 8711027), we followed the manufacturer’s instructions and maintained a constant dye concentration. In addition to PDAC cells, puromycin was included as a positive “kill-curve” control in most experiments to ensure measurement quality.

### Transmigration experiments

Endothelial cells were seeded onto fibronectin-coated coverslips (10 µg/mL) and allowed to reach confluence over 3-4 days. On the day of the experiment, lifeact-mNeonGreen or lifeact-mScarlet-I PDAC cells, resuspended in endothelial culture medium, were added to the endothelial cell monolayer. After 15 min, 2 h, or 8 h, samples were fixed with 4% paraformaldehyde (Thermo Fisher Scientific, 28908) in PBS or growth medium for 10 min at 37 °C. Importantly, no permeabilisation step was performed before immunostaining. Primary antibodies (1:200 dilution) were incubated for 1-2 h at room temperature, followed by incubation with species-appropriate secondary antibodies (1:400 dilution) for 45-90 min. Alexa Fluor 647 phalloidin (1:800) and DAPI (1:5000) were added alongside the secondary antibodies. Between each step, samples were washed with PBS. Coverslips were mounted using ProLong Glass mounting medium (Invitrogen, P36984). Imaging was performed with a Marianas spinning-disk microscope (20× air or 40x objectives). Maximum-intensity projected images were then processed to segment PDAC cells and fibronectin signals using automatic thresholding (default method) in Fiji (63). For PDAC cell morphometric analyses, segmented images were manually corrected, if needed, to separate individual cells before running particle analysis in Fiji.

### Live-cell imaging of transmigration

Ibidi 8-well slides were first coated with fibronectin (10 µg/mL) for 1 hour at 37 °C. After washing, 150,000 HUVECs expressing CAAX-EGFP (mosaic) were seeded and cultured for 3 days to form a confluent monolayer. Before imaging, fresh HEPES-containing medium (20 mM) was added, and the cells were transferred to a confocal Airyscan microscope (Zeiss, LSM880; see details in the dedicated section). PDAC cells (up to 100,000 cells per well), resuspended in HEPES-supplemented ECGM, were added to the HUVEC monolayer. Images were acquired in 3D at the fastest possible interval (ranging from 30 to 200 seconds, depending on FOV size and number of slices), with 15-25 z-slices at 0.17-0.2 µm z-steps. The images were processed with Airyscan in Zen Black (2.3), and 3D reconstructions were created with Arivis Vision4D (3.5.0).

Selection of imaging regions was based on two criteria for starting acquisition: a high density of CAAX-EGFP-expressing HUVECs and the presence of individual PDAC cell(s). We stopped acquisition after 1 h 30 min to 2 h if no signs of transmigration were identified (classified as “no transmigration”). Signs were variable and included PDAC protrusions, PDAC cell spreading, opening of endothelial cell-cell junctions, and endothelial retraction fibres. Retrospective analysis of the acquisitions allowed us to identify the first sign as the “event start”, and we defined the “end of transmigration” when the gap in the monolayer reached the area of the round PDAC cells at the start. Diameter and gap size were measured manually in Fiji using maximum projections. Classification of the transmigration mode was based on the presence of protrusions and contact between PDAC and endothelial cell membranes (“TC protrusion-based transmigration”); or the absence of long PDAC protrusions, separation between PDAC and HUVEC membranes, and the presence of retraction fibres from HUVEC cells (“EC retraction-based transmigration”). Scoring was not blinded, as PDAC cells are easily identifiable. Instead, ambiguous transmigration events showing signs of both categories were classified as “both mechanisms”.

### Light microscopy setup

The spinning-disk confocal microscope used was a Marianas spinning-disk imaging system with a Yokogawa CSU-W1 scanning unit mounted on an inverted Zeiss Axio Observer Z1 microscope, controlled by SlideBook 6 (Intelligent Imaging Innovations, Inc.). Images were acquired using an Orca Flash 4 sCMOS camera (2,048 × 2,048 pixels; Hamamatsu Photonics). The objectives used were 63× (NA 1.4 oil-immersion, Zeiss PLN Apo), 40x (NA 1.1 water, Zeiss LD C-Apochromat), 20× (NA 0.8 air, Zeiss Plan-Apochromat) and 10× (Plan-Apochromat 10×/0.45 NA Ph1).

Widefield imaging was performed using a Nikon Eclipse Ti2-E widefield microscope equipped with either a 10× Nikon CFI Plan-Fluor objective lens (NA 0.3) or a 20× Nikon CFI Plan Apo Lambda objective lens (NA 0.75).

The confocal microscope used was an LSM880 (Zeiss), equipped with an Airyscan detector (Carl Zeiss) and a 63× Zeiss C Plan-Apochromat objective (NA 1.4, oil immersion). The microscope was operated using Zen Black (2.3), with the Airyscan set to standard super-resolution mode.

The light-sheet microscope used was a ZEISS Lattice Light-sheet 7 (Carl Zeiss Microscopy GmbH, Jena, Germany). Samples were maintained at 37 °C and 5% CO_2_ throughout imaging. Samples were imaged every 10 minutes over several hours. For excitation and detection, a combination of 488/591/639 nm lasers, appropriate emission filters, and the Orca-Fusion sCMOS camera (Hamamatsu Photonics K.K.) was used with a dithered lattice light sheet measuring 15 µm × 0.55 µm. Images were acquired and processed (deskew and deconvolution) using ZEN (version 3.11).

### Scanning electron microscopy

Lifeact-mNeonGreen or Lifeact-mScarlet-I PDAC cells, resuspended in endothelial culture medium, were added to a confluent endothelial cell monolayer. At 15 min, 2 h, and 8 h, samples were fixed in PBS with 4% paraformaldehyde (Thermo Fisher Scientific, 28908) for 10 min at 37 °C. Increasing concentrations of EtOH baths (30%, 50%, 70%, 90%, and 100% ethanol, 3 min each) were used before adding hexamethyldisilazane (1:1 in EtOH 100%; 440191, Sigma-Aldrich) for 20 min; then the mixture was left to evaporate overnight. Dry coverslips were attached to aluminium stubs and sputter-coated with 2 nm of platinum. Imaging was performed with a Field-emission Scanning Electron Microscope (Thermo Scientific Apreo S) at an accelerating voltage of 2 kV. Events were classified, as explained in the previous section, based on PDAC protrusions, endothelial gaps, and retraction fibres from endothelial cells. Scoring was not blinded, as the PDAC cells are easily identifiable.

### Zebrafish model and injection with cancer cells

Adult zebrafish were maintained on a 12-hour light-dark cycle in stand-alone housing racks (Aqua-Schwartz, Göttingen, Germany). Embryos were collected via natural spawning in specialized 1.7 L sloped-bottom mating tanks (Tecniplast, Buguggiate, Italy). The *Tg(fli1:EGFP)* transgenic zebrafish line (ZFIN identifier y1Tg), in the Casper background and kindly provided by Dr Markus Affolter and Dr Heinz-Georg Belting (University of Basel, Switzerland), was used in these experiments. All zebrafish housing and experimental procedures were conducted under licences MMM/465/712-93 (Finnish Ministry of Agriculture and Forestry) and GTLK/004/E/2016 (Finnish Ministry of Social Affairs and Health), and within the Zebrafish Core Facilities at Turku Bioscience Centre.

Larvae were maintained in E3 medium (17.4 mM NaCl, 0.2 mM KCl, 0.1 mM MgSO_4_, 0.2 mM Ca(NO_3_)_2_), buffered with 0.15 mM HEPES (pH 7.6) and supplemented with 200 µM 1-phenyl-2-thiourea (Sigma-Aldrich, P7629) to prevent melanogenesis, as previously described (67). At 48 hours post-fertilisation (hpf), larvae were immobilised in 0.8% low-melting-point agarose and chemically anaesthetised in E3 medium supplemented with 650 µM tricaine (ethyl-3-aminobenzoate-methanesulfonate). PDAC cells were injected into the duct of Cuvier using a Nanoject microinjector 2 (Drummond) with micro-forged glass capillaries (25-30 µm inner diameter) filled with mineral oil (Sigma-Aldrich). Injections delivered 13 nL of a cell suspension at 100 × 10^6^ cells/mL. Larvae were positioned under an AxioZoom microscope (Zeiss) for injection (for more details, see (27)). After injection, embryos were kept in an agarose pad for 3 hours before imaging the caudal plexus region with a spinning-disk confocal microscope (3i CSU-W1) equipped with a 20× Zeiss Plan-Apochromat objective (NA 0.8). At the 24 hpi imaging time point, embryos were released from agarose after injection and remounted the next day for imaging on the same spinning-disk microscope. The *Tg(fli1:EGFP)* reporter allowed us to visualise vessel walls in 3D volumes and establish the relative positions of PDAC cells (intravascular or extravascular/extravasated).

For time-lapse imaging, embryos were kept on an agarose pad for 3 hours after injection, then imaged on either the same spinning-disk microscope or an Airyscan microscope (Zeiss) equipped with a 40x objective (NA 1.4). 3D rendering of the images was performed using IMARIS 10.4.

To generate preferred arrest-site heatmaps, 20× z-stacks were converted into maximum-intensity projections in Fiji (63). These projections were then affine-registered using the vascular reporter channel as the reference, with DRMIME (28). Tumour cells were segmented from the tumour fluorescence channel using StarDist2D (29). Both DRMIME and StarDist2D were implemented in ZeroCostDL4Mic and DL4MicEverywhere (68, 69). Pixel-wise cell density maps were calculated for each condition from the segmented cancer cells, normalised to the number of embryos, and lightly smoothed for heatmap display. Zebrafish larval anatomical sites were manually drawn on the heatmaps.

### Transwell assays

100,000 PDAC cells were cultured in their normal media on Transwell inserts (inside the well/upper compartment; Insert-Single 13 mm for 24-/6-well BRANDplates, BR782846, 3 µm pore size) deposited in 24-well plate wells for 2 days. In parallel, monolayers of endothelial cells were grown to confluence in 24-well plates as previously described. On the day of the experiment, ECGM media was changed in the endothelial cell wells, transwells were transferred to the endothelial cell wells, and the cells were left for 24 h. Fixation was performed after removing the transwell by incubating in 4% PFA for 10 min at 37 °C. FN-accessibility assay staining was performed, and the signal was quantified with Fiji (63). Fluorescently labelled PDAC cells were used in these experiments to ensure that no cells were in contact with the endothelial cell layer.

### Conditioned media

To generate conditioned media, we started with a 90%-confluent 10-cm dish of PDAC cells, then replaced the medium with the appropriate medium (10 mL of DMEM, RPMI, ECGM, basal ECGM, or ECGM from HUVEC culture) and incubated for 24 hours. Media were collected, filtered (0.45 µm Millipore, SLHVR33RS), and added to an Amicon® Ultra Centrifugal Filter (3 kDa MWCO, Millipore, UFC9003). The collected fraction ranged from 180 to 250 µL across conditions and repeats. These were aliquoted and stored at -80 °C, thawed only once, and used within 4 months. The working concentration in our experiments was 1:10 in ECGM, which can be estimated as a 5x concentration relative to the culture medium. For the FN-accessibility assay, the samples were fixed 2 hours after the addition of media. For impedance measurements, experiments ran for at least 15 hours after the addition of the media.

### Nearest-neighbour analysis of AsPC-1 cells and cleaved-caspase-3-positive endothelial cells

Confluent HUVEC, HP-MEC and HDMEC monolayers were incubated with Lifeact-mScarlet-I-expressing AsPC-1 cells for 8 h, fixed and stained with DAPI and an antibody against cleaved caspase-3. Twenty-five fields of view were randomly selected from each coverslip and acquired using a Marianas spinning-disk confocal microscope with a 20× objective and 2×2 camera binning.

Nearest-neighbour distances were measured using a custom Python pipeline based on NumPy, scikit-image and SciPy; Matplotlib was used for visualisation and quality control. Maximum-intensity projections of the z-stacks were analysed field by field. AsPC-1 cells were segmented from the Lifeact-mScarlet-I channel, and cleaved-caspase-3-positive objects were segmented by intensity thresholding using predefined lower and upper thresholds. Small objects were removed to suppress segmentation noise, and cleaved-caspase-3-positive objects smaller than 10 µm^2^ were excluded. To prevent apoptotic tumour cells from being misclassified as endothelial cells, cleaved-caspase-3-positive objects overlapping an AsPC-1 mask were excluded prior to object labelling and distance measurement.

For each segmented AsPC-1 cell, the centroid was calculated, and its Euclidean distance to the centroid of the nearest non-overlapping cleaved-caspase-3-positive object was measured. The nearest neighbour was defined as the cleaved-caspase-3-positive object with the shortest centroid-to-centroid distance. Annotated images showing object outlines and the corresponding nearest-neighbour connections were generated for quality control.

A spatially random reference distribution was generated separately for each endothelial cell type and biological replicate. For every field of view, point coordinates were sampled uniformly within the image boundaries and outside cleaved-caspase-3-positive masks. The number of random points was set to the mean number of segmented AsPC-1 cells per field of view in the corresponding endothelial cell type and biological replicate. The distance from each random point to the nearest cleaved-caspase-3-positive object in the same field of view was calculated using the same centroid-based procedure. Random points falling within cleaved-caspase-3-positive masks were excluded and resampled. Annotated control images were generated to verify point placement and nearest-neighbour assignment.

### Mouse models and injection with cancer cells

Female 9-to 15-week-old athymic nude mice were used in this study (Inotiv Netherlands, Hsd:Athymic Nude-Foxn1nu). Animals were housed in the common animal facility service UTU CAL in Turku. Animal experimentation was ethically assessed and authorised by the National Animal Experiment Board and was in accordance with the Finnish Act on Animal Experimentation (animal licence number ESAVI/6253/2024). Upon receipt, technical staff at the animal facility randomly allocated the animals to cages in numbers corresponding to our future experimental conditions (blindly to the experimenter). All efforts were made to minimise animal suffering and to reduce the number of animals used. Effectively, our longitudinal bioluminescence studies (15 min to 21 days) and the fluorescent immunostainings at different time points were performed on the same groups of animals. To achieve this, we injected a mix of PDAC cells expressing EGFP-Luciferase (2/3) for longitudinal study and Lifeact-mNeonGreen (1/3) for high-resolution imaging. This resulted in a progressive decrease in the number of animals per group and a loss of statistical power at the last time points in our experiments. This reduction in animal numbers was a trade-off we accepted because our scientific focus was on the earliest steps of metastatic seeding in the lungs. Animal numbers for the longitudinal bioluminescence and fluorescence analyses are provided in Table 1 and Table 2. Animal numbers for the saracatinib treatment experiment are provided in Table 3.

**Table 1.** Animal numbers used for the longitudinal bioluminescence analysis shown in Fig. 3C.

| Fig. 3C (n animals) | t0 | 6 h | 1 d | 2 d | 4 d | 7 d | 10 d | 14 d | 18 d | 21 d |
| --- | --- | --- | --- | --- | --- | --- | --- | --- | --- | --- |
| MIA PaCa-2 | 11 | 8 | 1 | 4 | 3 | 3 | 2 | 2 | 1 | 1 |
| AsPC-1 | 17 | 14 | 7 | 4 | 7 | 7 | 5 | 5 | 2 | 3 |
| Panc 10.05 | 9 | 7 | 1 | 4 | 3 | 3 | 2 | 2 | 1 | 1 |

**Table 2.** Animal numbers used for the lung fluorescence analysis shown in Fig. 3E.

| Fig. 3E (n animals) | t0 | 6 h | 1 d | 2 d | 7 d | 14 d | 21 d |
| --- | --- | --- | --- | --- | --- | --- | --- |
| MIA PaCa-2 | 3 | 2 | 1 | 0 | 0 | 1 | 0 |
| AsPC-1 | 3 | 3 | 3 | 1 | 1 | 2 | 2 |
| Panc 10.05 | 3 | 2 | 1 | 1 | 0 | 0 | 0 |

**Table 3.** Animal numbers used for the saracatinib treatment experiment shown in Fig. 6K-L.

| Fig. 6K-L (n animals) | t0 | 1 d | 2 d | 4 d | 7 d | 14 d | 21 d |
| --- | --- | --- | --- | --- | --- | --- | --- |
| Saracatinib | 19 | 15 | 10 | 10 | 9 | 9 | 9 |
| DMSO | 19 | 15 | 12 | 12 | 12 | 12 | 12 |

Several *ex vivo* imaging experiments were performed at the CRUK Scotland Institute. For those experiments, female Foxn1nu Nude (BALB/cAnNcrl) mice obtained from Charles River Laboratories (acclimatised for at least 1 week in our mouse facility) were used at 9 weeks of age. Mice were housed at the CRUK Scotland Institute, UK, in high-barrier conditions with a 12-hr light cycle at 19-22 °C and 45-65% humidity, with ad libitum access to food and water. Procedures were performed in accordance with the UK Animal (Scientific Procedures) Act 1986, approved by the local animal welfare committee (AWERB), and conducted under UK Home Office Licence (PP8411096) at the CRUK Scotland Institute.

On the day of the experiment, pancreatic ductal adenocarcinoma (PDAC) cells were resuspended at a concentration of 3 × 10^6^ cells/mL in PBS, and 100 µL of the cell suspension was administered by intravenous injection into the lateral tail vein. Mice were allowed to recover in their cage for 10 min. For bioluminescence imaging, D-luciferin was administered intraperitoneally (150 µL of 20 mg/mL solution in PBS). We waited 5 min for the luciferin to spread through the peritoneal cavity. Mice were transferred to the anaesthesia chamber (isoflurane, VETMEDIC, 2% at 1 L/min flow rate) of the imager (IVIS imaging system, Revvity), and imaging began 10 to 15 min after luciferin administration. Animals were positioned consistently with their nose in an isoflurane-dispensing mask and secured with adhesive tape during image acquisition. Bioluminescence images were acquired at low binning for 5 min of signal accumulation. The analysis ROI was standardised as a large square centred on the mouse lungs, and the results were normalised to the first acquisition time point for each animal. Following the imaging session, either the animals were allowed to recover in their home cage under surveillance, or, for animals that were to be sacrificed, they were transferred to the CO_2_ euthanasia chamber (see next paragraph). The remaining mice received repeated intraperitoneal injections of D-luciferin according to each protocol, at 3, 6, 9, or 24 hours, or 2, 3, 4, 7, 14, or 21 days after tumour cell injection. Mice were subsequently euthanised, and tissues were collected as described above.

Mice were euthanised by CO_2_ inhalation and pinned on a styrofoam board, ventral face up. The tracheal region was exposed, and a tracheotomy was performed. A silk suture (FST, 18020-01) was placed around the trachea without being tightened, and a small transverse incision (less than 1 mm) was made using spring scissors (FST, 15024-10). A 5 mL syringe fitted with a custom-made blunt-ended 21 G needle (Terumo, 050110C) was filled with 2 mL of low-melting-point agarose (Invitrogen, 16520050), maintained at 42-45°C to keep it liquid. The needle was gently inserted into the tracheal incision, and the silk suture was tightened around both the trachea and the needle to prevent leakage during inflation. The agarose solution was slowly injected until the lungs were fully inflated in the thoracic cage. Immediately after needle removal, the silk suture was tightened, and the lung was dissected (5 lobes in 1 piece). Lungs were immersed in ice-cold PBS and either processed for *ex vivo* analysis (see next section) or fixed in 4% PFA at 4°C overnight.

In case of intracardiac perfusion of fixative (Fig. 3D-G), the Harvard Apparatus Peristaltic Pump P-70 was used. 10 min of PBS perfusion followed by 10 min of 4% Paraformaldehyde perfusion were performed at a flow rate of 1ml/min, before dissection of the lungs and immersion in ice-cold PBS. In this case, no agarose inflation was performed.

Saracatinib treatment was done through gavage (Merck, CAD9917) at the concentration of 50 mg/kg/day for 5 days. The stock solution was prepared in DMSO at 180 mM. Gavage started the day before the injection, was given 4 hours before the injection on the day of injection, and lasted for 3 more days. Stock solution was prepared in 0.5% carboxymethyl cellulose in sterile PBS (1:15 dilution) and compared to the same concentration of carrier. Injection mix and procedures were as described previously. Schedule for bioluminescence measurements: 0, 1, 2, 4, 7, 14, 21 days. Sacrifice for lung inflation, dissection and immunolabeling: 9 hours post-injection to capture ongoing extravasation events.

### Tissue slice staining, imaging and quantifications

Dissected lungs were placed in ice-cold PBS before being sliced into 200 to 300 µm-thick sections on a vibrating microtome (5100mz; Campden Instruments or Leica VT1200) with stainless-steel blades (Model 7550-1-SS; Campden Instruments or Gillette platinum blade) at the highest oscillation amplitude and speed, advancing at 1 mm/s.

Tissue slices were first permeabilised and blocked with 10% Horse Serum (Gibco, 16050-122), 1% BSA (Sigma-Aldric, 0000454950), 0.3% Triton X-100 (Sigma-Aldrich, 1000882545), and Azide 0.05% (Sigma-Aldrich, 71289-5G) in PBS (permeabilisation and blocking solution) for 1 hour at room temperature (RT). Slices were then rinsed with a solution containing 1% BSA, 0.01% Triton X-100, and 0.05% Azide in PBS (rinsing/washing solution). Subsequently, slices were incubated overnight at 4 °C with primary antibody (1:200) in PBS supplemented with 10% Horse Serum, 1% BSA, 0.01% Triton X-100, and 0.05% Azide (antibody solution). After primary antibody incubation, slices were rinsed and washed three times for 20 minutes with the washing solution, then incubated with secondary antibodies (1:400) in the antibody solution for 3 hours at RT. Finally, slices were rinsed twice for 5 minutes each with the washing solution, rinsed with Milli-Q water, and mounted in ProLong™ Glass Antifade Mountant (Thermo Fisher Scientific, P36980). Mounted slices were incubated at RT overnight in the dark and then stored at 4 °C.

Tissue slices were imaged with a Marianas spinning-disk microscope using image tiling (low magnification), z-stacks, and multichannel excitation with 10×, 20×, 40×, and 63× objectives (see dedicated section). Low-magnification tiling images were segmented in Fiji (intensity-based on the green and DAPI channels). Large bronchial or venous areas were manually removed before using the DAPI segmentation (tissue area) to normalise the number of foci per mm^2^ of lung tissue. Analysed fields of view were randomly acquired via systematic x,y scanning of large lung slice areas until a sufficient number of fluorescent cancer cells were detected. Exclusion was decided by the experimenter only in cases of insufficient resolution due to the cancer cell being too far from the tissue and related light scattering. Cancer cells, endothelial cells, and the basal lamina were qualitatively assessed in three dimensions to classify phenotypes. Parameters included the continuity of the staining for collagen IV to characterise the formation of gaps, the appearance of strongly fluorescent large puncta that were distinguishable from the background signal of PECAM-1 staining, to characterise aggregates, and the localisation of cancer cells from both Collagen and PECAM to decide on intra- versus extravascular. Extravasation-associated vascular disruption was defined by the discontinuity and reorganisation of PECAM-1 and Collagen IV signals in 3 dimensions.

### Tumour cell segmentation from low-magnification immunostainings

For visualisation purposes, green fluorescent tumour cells were segmented from low-magnification mouse lung slice images using the ImageJ plugin Labkit (70). For each image, a separate pixel classifier was trained by manually annotating tumour cells visible against the autofluorescent background and the bronchiolar/large-vessel wall signal. The annotation and training process was repeated as many times as necessary to achieve satisfactory segmentation. The segmentation results were then exported, converted into Fiji masks, and overlaid on the original images to facilitate manual correction. The largest cancerous foci (AsPC-1, 21 days) were manually drawn. Particles smaller than 50 µm^2^ were filtered out to create the final masks, and the original images were then overlaid with these masks to produce the images shown in the figures.

### Precision-cut lung slices and ex vivo live imaging and analysis

The live PCLS procedure was performed as in (33). The tumour cells used were PDAC cells expressing Lifeact-mScarlet-I (described in the previous section). At the CRUK Scotland Institute, mice were humanely killed by overdose of anaesthetic (i.p. injection of sodium pentobarbital) followed by permanent cessation of circulation by severing the femoral artery. In Turku, CO_2_ inhalation was used. Tracheostomy, inflation and dissection were performed as described in the previous section. The largest lung lobe was isolated in PBS. After vibratome slicing (described above), one lung slice was stained with Alexa Fluor 488-conjugated anti-PECAM-1 (Mec13.3; 1:100; BioLegend, 102501) in complete medium (phenol-red free DMEM (Thermo Fisher Scientific, 21063029) supplemented with 10% FBS) for 20 minutes at 37 °C prior to imaging in a 4-well slide (ibidi, 80427) under imaging harps (Warner Instruments, SHD-22CL/15). The antibody was left in the well for the duration of the acquisition.

Slices were imaged on a Zeiss LSM880 Airyscan confocal microscope at 37 °C with 5% CO_2_. Lung slices were imaged for 10 to 20 hours with z-stacks, multichannel, and multipositioning every 15 minutes using Plan Apo objective lenses (20×/0.8 NA, Air; Carl Zeiss). Fields of view were randomly selected around cancer cells arrested in vessels.

Timewise, the mice were injected, left to recover for 5 minutes, sacrificed, and lung slices were prepared within 1 hour after sacrifice, except for the one imaged at 24 hours post-injection, which was left in the home cage for 24 hours before sacrifice. n=2 mice were used at the CRUK Scotland Institute, and n=8 mice were used in Turku, resulting in 45 acquired fields of view at 0 hours post-injection. n=1 mouse was used at the CRUK Scotland Institute at 24 hours post-injection, resulting in n=5 fields of view (not quantified). Airyscan 2D processing was performed on the raw acquisitions prior to channel realignment and registration using Fast4dReg (61). Cancer and endothelial cell behaviours were qualitatively assessed in three dimensions to classify phenotypes: tumour cell death, endothelial retraction, cancer cell protrusions, uncharacterized extravasation (for ambiguous observations), and intravascular and extravasated cells. Selected acquisitions were optimised in ImageJ for visualisation (z-projections, time-point isolation, adjusted LUT, and inverted LUT).

## Statistical analysis

The numerical data used for the figures have been archived on Zenodo (30). Statistical analyses are detailed in the figure legends. Where applicable, normality was assessed using the Shapiro–Wilk test. Quantitative analyses included Welch’s *t*-test, the Mann–Whitney U test, one-way ANOVA, and the Kruskal–Wallis test followed by Dunn’s or Holm–Šídák post hoc test, as specified in the corresponding figure legend. Categorical data were analysed using the chi-squared (χ^2^) test.

## Data availability

The raw numerical values and images used to build this manuscript have been archived on Zenodo (30). Any additional information required to reanalyse the data reported in this paper is available from the corresponding authors.

## Manuscript preparation

Figures were prepared using Adobe Illustrator 2024 and CS6. Data were plotted using Microsoft Excel and GraphPad Prism v10 and v11. GPT-5.5 (OpenAI) and Grammarly (Grammarly, Inc.) were used as writing aids while preparing this manuscript. The authors further edited and validated all sections of the text. The preprint version of this manuscript was formatted with Rxiv-Maker (71).

## Author contributions

**Conceptualisation**: G.F., J.I., G.J. **Methodology**: G.F., J.W.P., A.I., M.R.C., F.F., E.M., J.R.W.C., M.S., S.A.W., L.M.C., J.I., G.J. **Reagents**: G.F., M.D.D., E.M., M.S., S.A.W., L.M.C. **Formal Analysis**: G.F., S.G., M.V., J.W.P., A.I., M.R.C., G.J. **Investigation**: G.F., S.G., M.V., O.J., H.H., G.J. **Writing - Original Draft**: G.F., J.I., G.J. **Writing - Review and Editing**: Everyone. **Visualisation**: G.F., S.G., H.H., J.W.P., A.I., M.R.C., G.J. **Supervision**: G.F.,, G.J. **Funding Acquisition**: G.F., J.I., G.J.

## Acknowledgments

This study was funded by the Research Council of Finland (338537, 371287, and 374180 to G.J., 360775 to J.R.W.C., 332402 to G.F., 346131 and 364182 to S.A.W. and J.I., and Academy Professorship 369992 to J.I.), the Sigrid Juselius Foundation (to J.I. and G.J.), the Cancer Foundation Finland (Syöpäjärjestöt; to G.J., J.I., and M.S.), the Jane and Aatos Erkko Foundation (to J.I.), and the Solutions for Health strategic funding for Åbo Akademi University (to G.J.). This research was also supported by the InFLAMES Flagships Programme of the Research Council of Finland (decision numbers: 337530, 337531, 357910, and 357911). G.J. is supported by the Finnish Cancer Institute (K. Albin Johansson Professorship). G.F. was supported by the Turku Collegium for Science, Medicine, and Technologies. M.D.D., F.F., and L.M.C. were funded by CRUK SI core programme award A23983, core funding to the CRUK Scotland Institute A31287, and the CRUK Scotland Centre CTRQQR-202100006. F.F. and L.M.C. were funded by Breast Cancer Now (2019DecPR1424). S.A.W. was supported by the Max Planck Society. This project was supported by an ERC grant (BorderControl; grant agreement number 101142305 to J.I.). Funded by the European Union. Views and opinions expressed are, however, those of the author(s) only and do not necessarily reflect those of the European Union or the European Research Council Executive Agency. Neither the European Union nor the granting authority can be held responsible for them.

Imaging was performed at the Advanced Imaging Core Facility at the Turku Bioscience Centre and at the Material Research Infrastructure (MARI), with samples prepared at the Electron Microscopy Core Facility at the Institute of Biomedicine, University of Turku. These facilities received support from Turku BioImaging, Biocenter Finland, and the Finnish Advanced Microscopy Node of Euro-BioImaging Finland (funded by the Research Council of Finland, FIRI 2023 grant decision numbers 359073 and 358879, and FIRI 2024 grant decision numbers 367582 and 367577). The authors thank the BioOpticService unit at the Max Planck Institute for Molecular Biomedicine for access to imaging instrumentation and technical support during a research visit, as well as the Beatson Advanced Imaging Resource (BAIR; RRID: SCR_023875) at the CRUK Scotland Institute for similar support during a research visit. The authors also thank the staff of the University of Turku Central Animal Laboratory, especially Joonas Khabbal and Emra Yatkin, for experimental assistance. Testament funds from Henna Ruusunen also supported this work.

## ABOUT THIS MANUSCRIPT

This work is licensed under CC BY 4.0.

## Supplementary Information

***Supplementary Movie 1: Transendothelial migration of pancreatic cancer cells through a HUVEC monolayer***. Panc 10.05 (Part A), MIA PaCa-2 (Part B), and AsPC-1 (Part C) cells expressing Lifeact-mScarlet-I were seeded onto HUVEC monolayers with mosaic expression of the membrane marker CAAX-EGFP and imaged by Airyscan confocal microscopy. A HUVEC monolayer without cancer cells is shown as a control (Part D). Maximum-intensity projections and 3D reconstructions are shown, with pauses and arrows highlighting key events. 3D reconstructions were generated using Arivis Vision4D (3.5.0).

***Supplementary Movie 2: MIA PaCa-2 cell transmigration through a HUVEC monolayer into a three-dimensional extracellular matrix***. MIA PaCa-2 cells expressing Lifeact-mScarlet-I were added to HUVEC monolayers with mosaic expression of the membrane marker CAAX-EGFP, grown on a collagen gel, and imaged by lattice light-sheet microscopy. Time-lapse image sequences and 3D reconstructions are shown. The 3D reconstructions were generated using IMARIS.

***Supplementary Movie 3: Pancreatic cancer cell extravasation in zebrafish embryos***. MIA PaCa-2 (Part A) and AsPC-1 (Part B) cells expressing Lifeact-mScarlet-I were injected into the duct of Cuvier of Fli1a transgenic zebrafish embryos expressing eGFP in endothelial cells, and the embryos were imaged by spinning-disk confocal microscopy. Each video begins with a maximum-intensity projection of a multi-channel overview of a larger FOV, followed by magnified multi-channel ROIs shown alongside the corresponding endothelial channel in grayscale. Key events are highlighted with yellow arrows.

***Supplementary Movie 4: AsPC-1 cells in fixed mouse lung slices***. AsPC-1 cells expressing EGFP-luciferase were injected into the tail vein of nude mice. Six hours after injection, the mice were sacrificed, and the lungs were dissected, sectioned, immunofluorescence-stained, and imaged with a spinning-disk confocal microscope. In Part A, a four-channel 3D image stack is shown alongside the corresponding PECAM-1 and collagen IV channels for the ROI. Similarly, in Part B, a four-channel 3D image stack is shown alongside the corresponding AsPC-1 and cleaved caspase-3 channels for the ROI.

***Supplementary Movie 5: AsPC-1 cell extravasation associated with endothelial retraction in mouse lung slices***. AsPC-1 cells expressing EGFP-luciferase were injected into nude mice via the tail vein. One hour post-injection, the mice were sacrificed, and the lungs were dissected, sectioned, immunolabelled for PECAM-1, and imaged live on an Airyscan confocal microscope. Maximum-intensity projection videos of a single AsPC-1 cell (Part A), a cluster of AsPC-1 cells (Part B), and control mouse endothelium without AsPC-1 cells (Part C) are shown. Videos containing cancer cells are presented as multi-channel videos, with the corresponding endothelial channel displayed in grayscale. For each video, the endothelial channel from the first and last frames is shown side by side in grayscale at the end of the video. Key events are indicated with arrows.

***Supplementary Movie 6: Additional examples of AsPC-1 cell extravasation associated with endothelial retraction in mouse lung slices*** . AsPC-1 cells expressing EGFP-luciferase were injected into nude mice via the tail vein. Approximately 1 hour post-injection, the mice were sacrificed, and the lungs were dissected, sectioned, immunolabelled for PECAM-1, and imaged live on an Airyscan confocal microscope. Maximum-intensity projection videos of two representative examples (Parts A and B) of AsPC-1 cell extravasation are shown. Each video is presented as a multichannel video alongside the corresponding endothelial channel in greyscale. For each video, the endothelial channel from the first and last frames is shown side by side in greyscale at the end of the video. Key events are indicated with arrows.

***Supplementary Movie 7: Different fates of AsPC-1 cells in mouse lung slices***. AsPC-1 cells expressing EGFP-luciferase were injected into nude mice via the tail vein. Approximately 1 hour post-injection, the mice were sacrificed, and the lungs were dissected, sectioned, immunolabelled for PECAM-1, and imaged live using an Airyscan confocal microscope. Maximum-intensity projection videos of AsPC-1 cell death within the vasculature (Part A) and prolonged intravascular retention without disrupting the mouse lung endothelium (Part B) are shown. Part A shows a representative example of AsPC-1 cell death, presented as a multichannel video alongside the corresponding cancer cell channel in greyscale. Part B shows a representative example of an AsPC-1 cell remaining within the vasculature, presented as a multichannel video alongside the corresponding endothelial channel in greyscale.

***Supplementary Movie 8: AsPC-1 cell migration in the mouse lung stroma 24 hours post-injection***. AsPC-1 cells expressing EGFP-luciferase were injected into the tail vein of nude mice. Twenty-four hours post-injection, the mice were sacrificed, and the lungs were dissected, sectioned, immunolabelled for PECAM-1, and live-imaged on an Airyscan confocal microscope. A maximum-intensity projection of a multi-channel video is shown alongside the corresponding endothelial channel in greyscale.

**Sup. Fig. S1.**
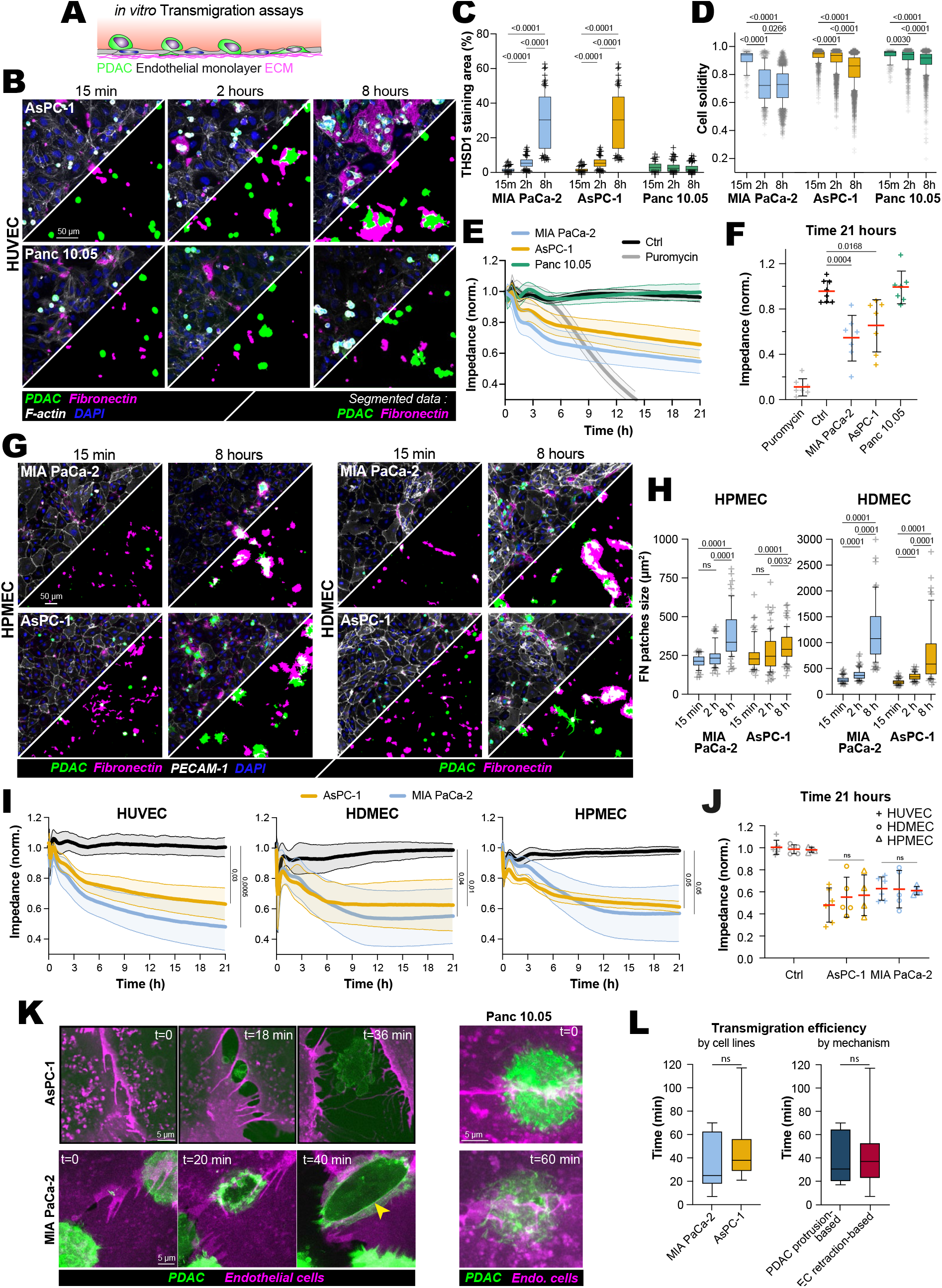
PDAC-induced endothelial barrier disruption is cell line dependent and conserved across different endothelia. (**A**) Schematic representation of the assay used to study PDAC cell arrest and transmigration across endothelial monolayers. (**B-D**) PDAC cells expressing Lifeact-mNeonGreen were added to HUVEC monolayers and fixed after 15 min, 2 h, or 8 h. Samples were stained without permeabilisation using DAPI, phalloidin, and anti-fibronectin or anti-thrombospondin-1 antibodies, and imaged by spinning-disk confocal microscopy. Cells and fibronectin-covered areas in the images were then segmented and quantified. (**B**) Representative images of AsPC-1 and Panc 10.05 cells, with corresponding cancer cell and fibronectin segmentation masks. Scale bar: 50 µm. (**C**) Quantification of accessible thrombospondin-1, expressed as percentage coverage per field of view, for each cell line and time point (n > 80 fields of view; 3 biological replicates). (**D**) Quantification of the solidity of segmented single-cell masks for each cell line and time point (n > 169 cells; 3 biological replicates). (**E-F**) PDAC cells were plated onto confluent HUVEC monolayers, and endothelial barrier integrity was monitored using xCELLigence impedance measurements. (**E**) Impedance traces were recorded for 21 h after PDAC cell addition. Data were normalised to the time of PDAC cell addition (t = 0 h). Puromycin (1 µg/mL) was used as a positive control for barrier collapse (n > 7 biological replicates). (**F**) Endpoint impedance values at t = 21 h, plotted as mean ± SD with individual data points (n > 7 biological replicates). P-values were determined using a Kruskal-Wallis test followed by Dunn’s post-test. (**G, H**) MIA PaCa-2 and AsPC-1 cells expressing Lifeact-mNeonGreen were added on top of HPMEC and HDMEC monolayers and fixed after 15 min, 2 h, or 8 h. Samples were stained without permeabilisation using DAPI, anti-PECAM-1, and anti-fibronectin, and imaged by spinning-disk confocal microscopy. (**G**) Representative images, together with corresponding cancer cell and fibronectin segmentation masks, are shown. Scale bar: 50 µm. (**H**) Quantification of accessible fibronectin (average patch size per field of view) for each cell line and time point (n > 55 fields of view; 3 biological replicates). (**I-J**) PDAC cells were plated onto confluent HUVEC, HPMEC and HDMEC monolayers, and endothelial barrier integrity was monitored using xCELLigence impedance measurements. (**I**) Impedance traces were recorded for 21 h after PDAC cell addition. Data were normalised to the time of PDAC cell addition (t = 0 h) (n > 4 biological replicates). (**J**) Endpoint impedance values at t = 21 h, plotted as mean ± SD with individual data points (n > 4 biological replicates). P-values were determined using a one-way ANOVA comparing each triplet of endothelial cell conditions on the x-axis, omitting comparisons involving PDAC conditions (as shown in panel **I**). (**K, L**) PDAC cells expressing Lifeact-mScarlet-I were added on top of a HUVEC endothelial monolayer expressing CAAX-EGFP (mosaic expression) and imaged using an Airyscan confocal microscope (related to Fig. 1E). (**K**) Representative still images extracted from live-cell microscopy recordings of Panc 10.05, AsPC-1, and MIA PaCa-2 cells. The yellow arrow indicates a contractile ring around an MIA PaCa-2 cell. Scale bar: 5 µm. (**L**) Additional quantification of transmigration dynamics from the live-cell imaging dataset shown in Fig. 1E-H. Left: time required to complete transmigration between MIA PaCa-2 and AsPC-1 cells. n = 7 events for MIA PaCa-2 and n = 9 events for AsPC-1. Right: time required to complete transmigration, grouped by transmigration mode. n = 8 events per mode. TC, tumour cells; EC, endothelial cells. (**C, D, H, L**) Results are presented as boxplots, with whiskers extending from the 10th to the 90th percentiles. The boxes capture the interquartile range, with the median marked by a line within each box. Data points outside the whiskers are depicted as individual crosses. P-values were determined using a Kruskal-Wallis test followed by Dunn’s post-test. The raw numerical values and images used to make this figure have been archived on Zenodo @follain2026Zenodo.

**Sup. Fig. S2.**
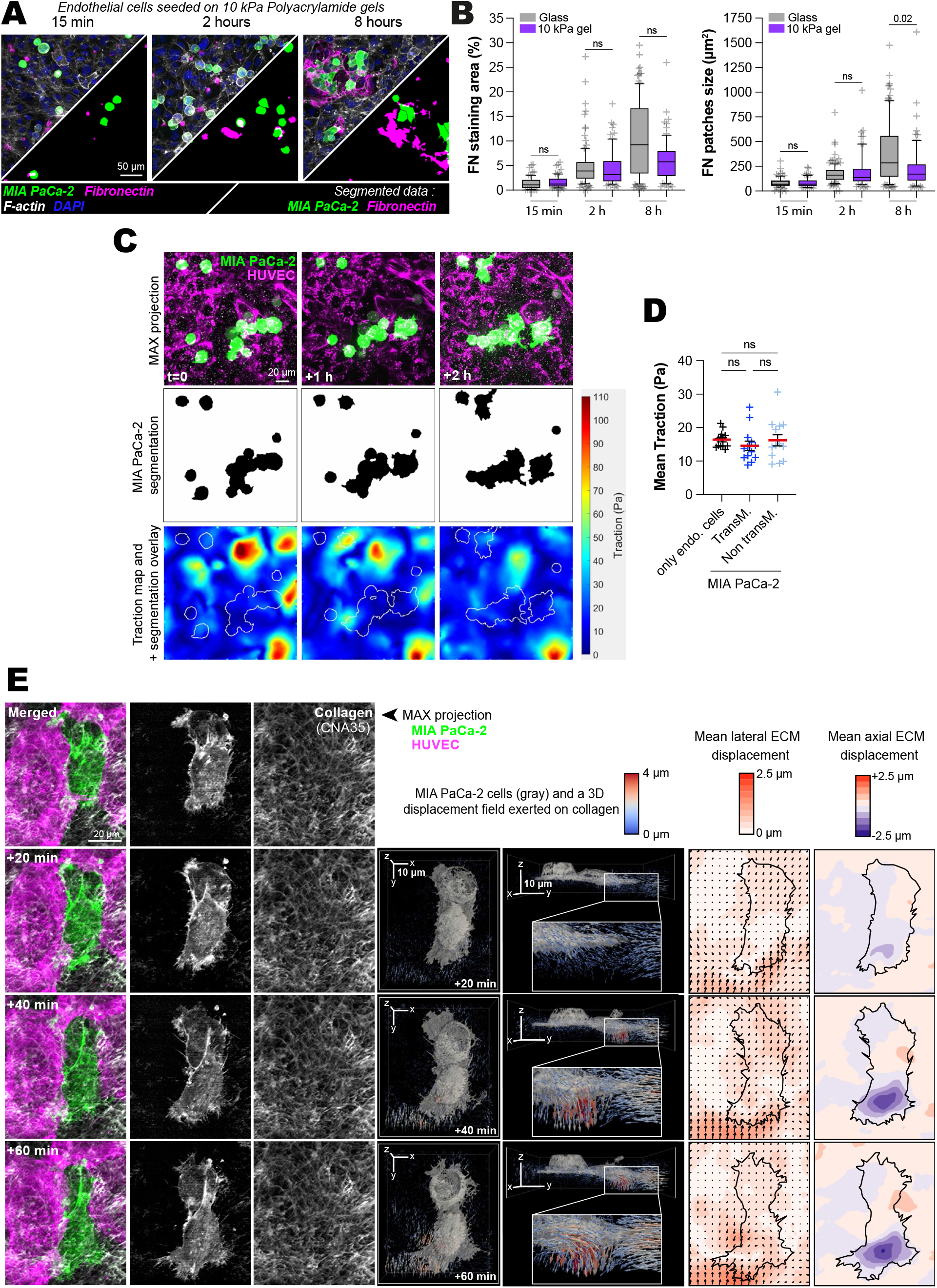
PDAC-mediated endothelial barrier disruption persists on compliant and three-dimensional matrices. (**A-B**) MIA PaCa-2 cells expressing Lifeact-mNeonGreen were added to HUVEC monolayers grown on 10 kPa polyacrylamide gels and fixed after 15 min, 2 h, or 8 h. Samples were stained without permeabilisation with DAPI, phalloidin, and an anti-fibronectin antibody and imaged using a spinning-disk confocal microscope. (**A**) Representative images and corresponding segmentation masks used for quantification. Scale bar: 50 µm. (**B**) Quantification of accessible fibronectin, expressed as percentage coverage and mean patch size per field of view, comparing endothelial monolayers grown on glass or 10 kPa gels at each time point (n > 70 fields of view; 3 biological replicates). Data are shown as boxplots. Boxes indicate the interquartile range, centre lines indicate the median, whiskers extend from the 10th to the 90th percentiles, and points outside the whiskers are shown as individual crosses. P values were determined using a Kruskal-Wallis test followed by Dunn’s post-test. Statistical comparisons are shown only for glass versus gel at each time point. (**C-D**) MIA PaCa-2 cells were added onto HUVEC monolayers grown on fluorescent bead-embedded polyacrylamide gels, and transmigration events were recorded by live-cell imaging using a spinning-disk confocal microscope. Cells were then removed with SDS, and beads were imaged again for 2D traction force microscopy analysis. MIA PaCa-2 masks generated by intensity-based segmentation were dilated to define regions potentially affected by tumour cells. These regions were then manually classified as transmigrating MIA PaCa-2 cells, non-transmigrating apically adherent MIA PaCa-2 cells, or endothelial-cell-only regions. (**C**) Representative images illustrating the experimental setup and analysis pipeline. (**D**) Quantification of mean lateral traction force per annotated region. n = 13 fields of view per condition. Data are shown as individual fields of view with mean ± SD. P values were determined using a Kruskal-Wallis test followed by Dunn’s post-test. (**E**) Live lattice light-sheet acquisition of transmigrating MIA PaCa-2 cells through a layer of endothelial cells grown on a collagen and fibronectin gel, analysed to determine the lateral and axial displacement of the collagen matrix during the process. From left to right: Merged channels of the acquisition (z-projection after deskewing), single tumour cell channel, single collagen channel, top and side views of 3D segmentation of the tumour cells overlaid with collagen displacement fields, mean lateral collagen displacements, and mean axial collagen displacements (see method details). The raw numerical values and images used to make this figure have been archived on Zenodo @follain2026Zenodo.

**Sup. Fig. S3.**
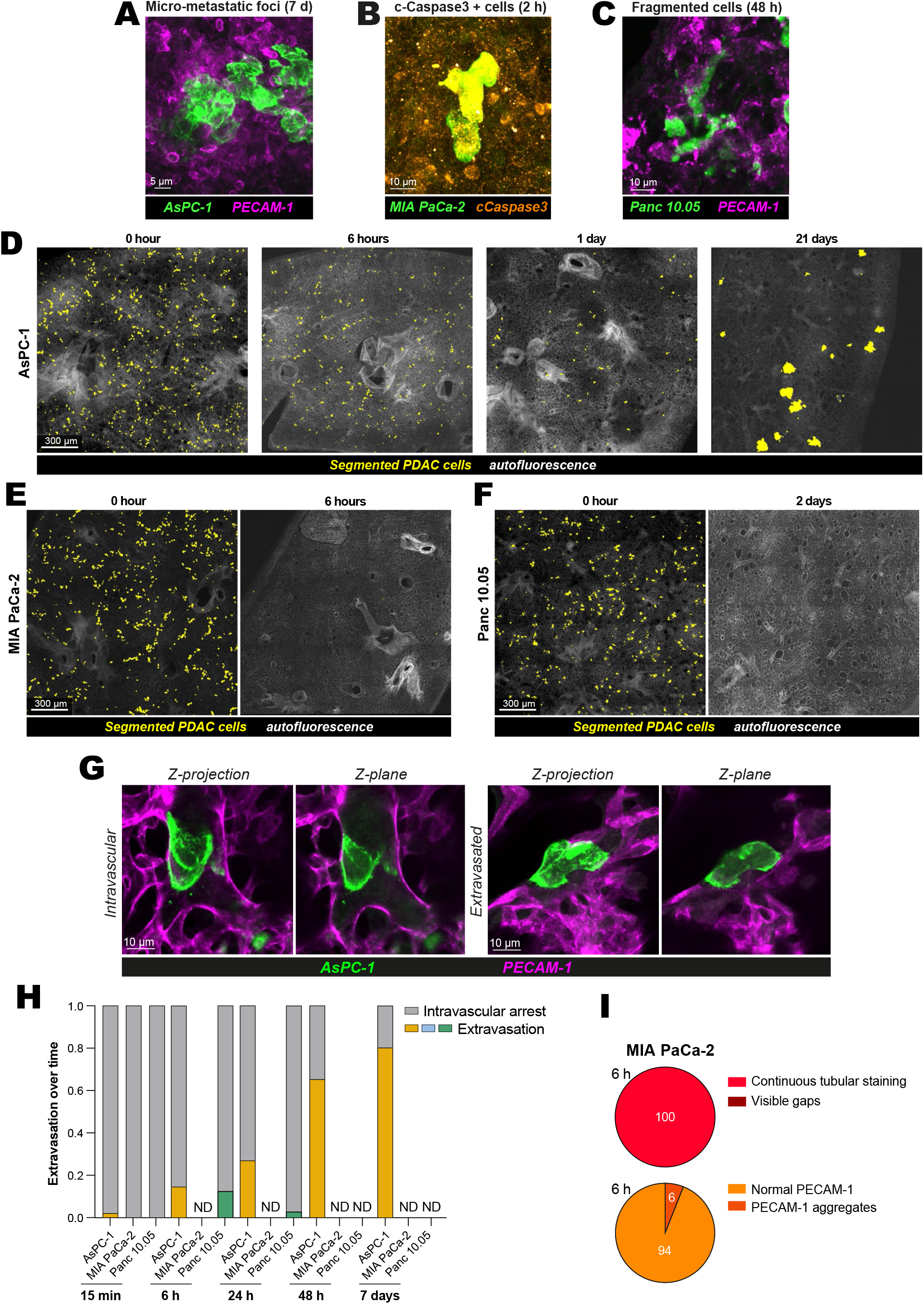
Differential lung retention and extravasation of PDAC cell lines after tail-vein injection in nude mice. (**A**) High-resolution confocal z-projection of a growing micrometastatic lesion representative of AsPC-1 behaviour 7 days post-injection (related to Fig. 3D). Scale bar: 5 µm. (**B**) High-resolution confocal z-projection of a cluster of dying MIA PaCa-2 cells labelled for cleaved caspase-3 at 2 h post-injection. Scale bar: 10 µm. (**C**) High-resolution confocal z-projection of fragmented Panc 10.05 cells, mostly intravascular, representative of Panc 10.05 behaviour 2 days post-injection. Scale bar: 10 µm. (**D-F**) Low-magnification confocal z-projections corresponding to Fig. 3E and showing key time points for each cell line. Initial lung lodging was similar across the three cell lines. (**D**) AsPC-1 time course showing an initial decrease in the number of foci, followed by outgrowth of a subset of foci. (**E**) MIA PaCa-2 time course, showing near-complete disappearance within the first hours post-injection. (**F**) Panc 10.05 time course, showing a slower disappearance than MIA PaCa-2. Scale bar: 300 µm. (**G**) Representative z-projection and selected optical plane showing examples of intravascular and extravasated AsPC-1 cells. AsPC-1 cells express Lifeact-mNeonGreen, and the endothelium is labelled by PECAM-1 immunostaining. Scale bar: 10 µm. (**H**) Quantification of extravasation related to (**G**). A comparable amount of biological material was immunostained and manually scanned for PDAC cells at each time point and for each cell line. Number of foci analysed, from left to right: n = 256, 173, 143, 158, 16, 121, 134, 37, 89, 2, 273, and 2. Groups with n < 40 were considered non-discriminant (ND) because of insufficient sample size. (**I**) Quantification of Collagen IV gaps and PECAM-1 aggregation around MIA PaCa-2 tumour cells using high-resolution z-stacks from the same lung slices as in Fig. 3D. Numbers of foci analysed, from left to right: n = 40 and 49. The raw numerical values and images used to make this figure have been archived on Zenodo @follain2026Zenodo.

**Sup. Fig. S4.**
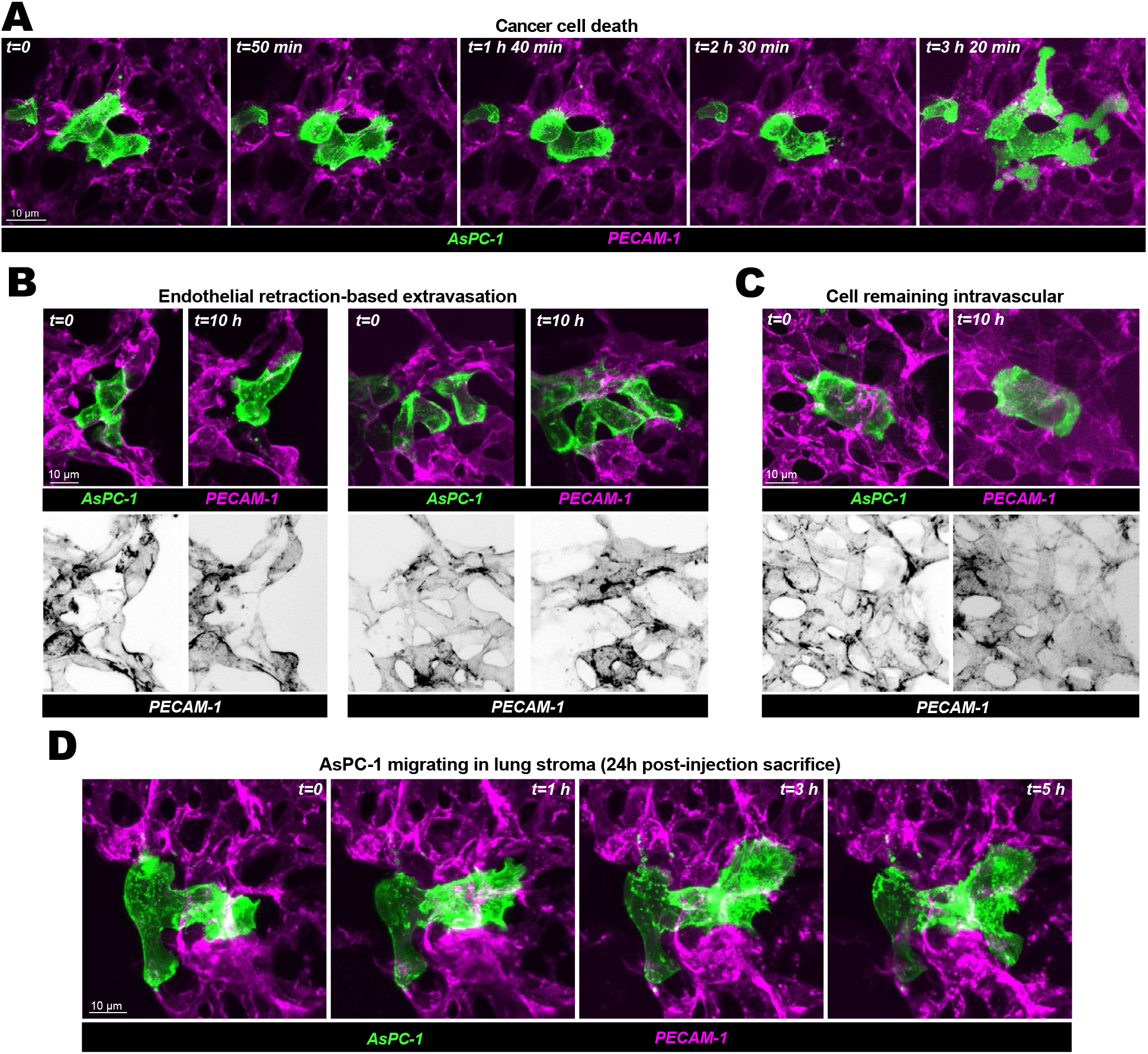
Live ex vivo imaging of AsPC-1 behaviour in lung slices. (**A-D**) Lifeact-mNeonGreen-expressing AsPC-1 cells were injected into nude mice via the tail vein. Mice were sacrificed either shortly after cancer cell injection (**A-C**) or 24 h post-injection (**D**). Lungs were then extracted, thick-sectioned, labelled for PECAM-1, and imaged ex vivo. (**A-C**) Time series extracted from live *ex vivo* lung-slice imaging experiments, related to Fig. 3H-L. (**A**) Example of an AsPC-1 cell dying within the vasculature. (**B**) Two examples of AsPC-1 extravasation associated with local endothelial retraction. (**C**) Example of an AsPC-1 cell remaining intravascular during the imaging period. Scale bars: 10 µm. (**D**) Time series extracted from a live ex vivo experiment started 24 h post-injection to capture the behaviour of AsPC-1 cells after extravasation. Cells are migrating in the lung tissue. Scale bar: 10 µm. The raw numerical values and images used to generate this figure have been archived on Zenodo (30).

**Sup. Fig. S5.**
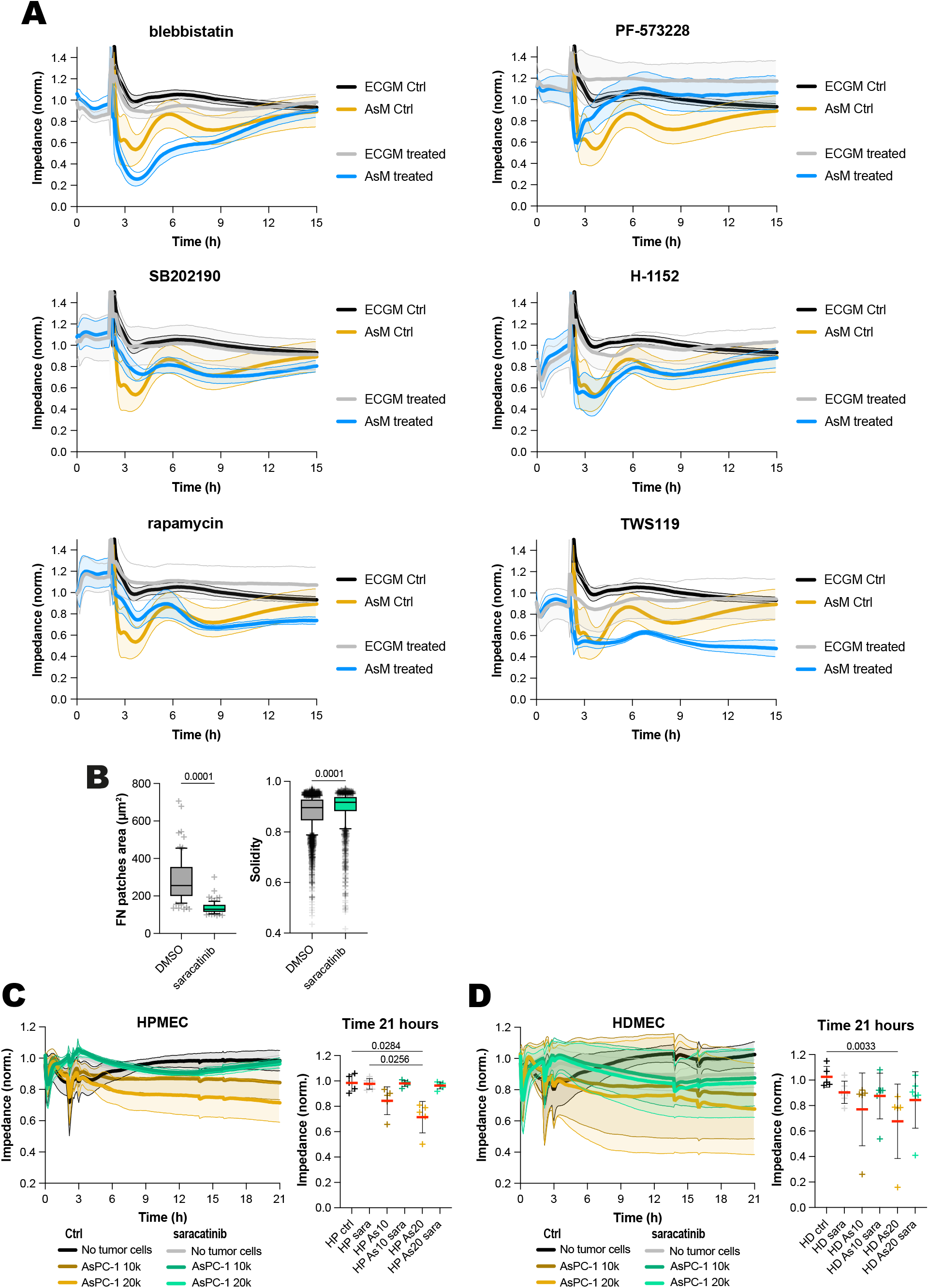
Saracatinib protects endothelial barriers from AsPC-1-induced destabilisation. (**A**) Complete impedance traces from the inhibitor screen shown in Fig. 6A. HUVEC monolayers were pre-treated with the indicated inhibitors for 2 h before addition of AsM-conditioned medium, and impedance was recorded over time. n > 2 biological repeats. (**B**) Additional quantification from the fibronectin-accessibility assay shown in Fig. 6C-D, comparing AsPC-1-induced endothelial barrier disruption in the presence or absence of saracatinib (n > 75 fields of view and n > 2222 cells; 3 biological replicates). (**C-D**) Impedance measurements of HPMEC (**C**) and HDMEC (**D**) monolayers treated with saracatinib and challenged with AsPC-1 cells. Impedance traces were recorded over 21 h. Endpoint impedance values at t = 21 h were plotted as mean ± SD with individual data points. n = 5 biological replicates. (**B, C, D**) P values were determined using a Kruskal-Wallis test followed by Dunn’s post-test. Raw numerical values and images used to generate this figure have been archived on Zenodo (30).

